# Reprogramming of lactate metabolism is linked to the oncogenesis of the virus-induced leukemia

**DOI:** 10.1101/2022.08.12.503711

**Authors:** Kosuke Toyoda, Jun-ichirou Yasunaga, Yuichiro Arima, Kenichi Tsujita, Azusa Tanaka, Osama Hussein, Miyu Sonoda, Miho Watanabe, Takafumi Shichijo, Daisuke Kurita, Kazutaka Nakashima, Kyohei Yamada, Hiroaki Miyoshi, Koichi Ohshima, Masao Matsuoka

## Abstract

Acceleration of glycolysis is a common trait of cancer. A key metabolite, lactate, is properly secreted from cancer cells, since its accumulation is toxic. Here, we report that a viral oncogene, HTLV-1 bZIP factor (HBZ), bimodally upregulates TAp73 to promote lactate excretion from adult T-cell leukemia-lymphoma (ATL) cells. HBZ protein binds to EZH2 and reduces its occupancy of the TAp73 promoter. Meanwhile, *HBZ* RNA activates TAp73 transcription via the BATF3-IRF4 machinery. TAp73 upregulates the lactate transporters MCT1 and MCT4. Inactivation of TAp73 leads to intracellular accumulation of lactate, inducing cell death in ATL cells. Furthermore, TAp73 knockout diminished development of inflammation in HBZ-transgenic mice. An MCT1/4 inhibitor, syrosingopine, decreased the growth of ATL cells in vitro and in vivo. MCT1/4 expression was positively correlated with TAp73 in many cancers, and their upregulations were associated with dismal prognosis. Activation of the TAp73-MCT1/4 pathway could be a common mechanism contributing to oncogenesis.

## Introduction

The core capabilities in the Hallmarks of Cancer, a framework for understanding the common features of diverse human malignant tumors, include “deregulating cellular metabolism” and “epigenetic reprogramming” (Hanahan, 2022; Hanahan and Weinberg, 2011). The Warburg effect is a well-known energy-generation phenomenon associated with cancer metabolism, by which the rate of glucose uptake, glycolysis, and consequent production of lactate are drastically increased in cancer cells even under aerobic conditions (Koppenol et al., 2011; Warburg, 1956). Epigenetic alterations, such as CpG DNA hyper- or hypomethylation and aberrant histone modifications, dysregulate chromatin structure and gene expression to promote oncogenesis. Although many types of malignancy share these hallmarks and enabling characteristics, the pathogenesis of each tumor is complex. To elucidate the molecular mechanisms for oncogenesis of each malignant disease, one must to consider the characters of each cell type and the associated exogenous factors such as pathogens.

Human T-cell leukemia virus type 1 (HTLV-1) is an oncogenic retrovirus that mainly infects CD4^+^ T cells and causes a fatal malignancy called adult T-cell leukemia-lymphoma (ATL). Since viral replication of HTLV-1 is generally suppressed *in vivo*, persistent infection is established mainly by clonal proliferation of infected cells. Among the viral genes, only *HTLV-1 bZIP factor (HBZ)*, which is encoded in the minus strand of the provirus, is conserved and expressed in all ATL cases. Knockdown of *HBZ* reduces the proliferation of ATL cell lines (Satou et al., 2006), and transgenic mice that express *HBZ* in CD4^+^ T cells (HBZ-Tg mice) develop systemic inflammation and T-cell lymphoma (Satou et al., 2011), suggesting that *HBZ* is significant in the pathogenesis of HTLV-1. A fascinating feature of the *HBZ* gene is that its transcript not only encodes the HBZ protein but also acts like long non-coding RNA (lncRNA) (Toyoda and Matsuoka, 2022). Previous studies using microarray analysis suggested that *HBZ* RNA promotes expansion of CD4^+^ T cells by inducing cell cycle- and apoptosis-related genes (Mitobe et al., 2015; Satou *et al*., 2006). Meanwhile, HBZ protein modulates the immunophenotypes of expressing cells toward a regulatory T cell (Treg)-like phenotype by inducing Treg-associated genes such as *Foxp3* (Kinosada et al., 2017; Yasuma et al., 2016; Zhao et al., 2011). Thus, *HBZ* plays critical roles in the clonal expansion of infected cells and the development of ATL by using both its coding and noncoding functions.

In this study, we show that *HBZ* induces reprogramming of glucose metabolism by upregulating the cellular transcription factor TAp73, and that HBZ also modulates the function of the enhancer of zeste homolog 2 protein (EZH2), leading to widespread epigenetic alterations. *HBZ* RNA and HBZ protein each activate the promoter of *TAp73* by different mechanisms, and TAp73 consequently induces MCT1 and MCT4, which act as lactate exporters to optimally excrete lactate created in ATL cells. EZH2 is one of the subunits of polycomb repressive complex 2 (PRC2), which functions as a methyltransferase to repress transcription by trimethylating histone H3 at lysine 27 (H3K27me3) (Laugesen and Helin, 2014). Gain-of-function mutations of EZH2 are found in several cancer types, whereas ATL is shown to have high expression levels of unmutated EZH2 (Fujikawa et al., 2016; Sasaki et al., 2011; Yamagishi et al., 2012). Here we show that HBZ protein binds to EZH2 and changes the genome-wide distribution of H3K27me3, resulting in high expression of TAp73 as well as other widespread changes to the transcriptome. In addition, TAp73 further activates transcription of *EZH2* in ATL cells.

## Results

### Both HBZ protein and RNA upregulate TP73

Both HBZ protein and *HBZ* RNA modulates cellular transcriptome through dysregulation of the promoter activity of the target genes (Ma et al., 2021; Matsuoka and Jeang, 2007; Matsuoka and Mesnard, 2020; Mitobe *et al*., 2015). To comprehensively analyze the target genes of HBZ protein and RNA, we began by conducting RNA sequencing (RNA-seq) and assay for transposase-accessible chromatin sequencing (ATAC-seq) of primary murine CD4^+^ T cells expressing wild-type *HBZ,* mutant *HBZ* that can act only in protein form, or mutant *HBZ* that can act only in RNA form (Figure 1A). Although the modes of action of HBZ protein and RNA were expected to be different, we found that all molecular forms of *HBZ* (WT, protein mutant and RNA mutant) had similar transcriptional profiles in some gene clusters (Figure 1B). Comparative analysis showed that 467 upregulated genes (Figure 1C, left) and 371 altered opened-chromatin regions (Figure 1C, right) were affected by all forms of *HBZ*. Indeed, *Ccr4* and *Tigit* are already known to be common targets, and the present results confirmed this to be the case (Figure 1D) (Kinosada *et al*., 2017; Sugata et al., 2016; Yasuma *et al*., 2016). Gene set enrichment analysis (GSEA) also supported the observation that both HBZ protein and *HBZ* RNA target identical gene sets, such as TP53- or KRAS-related gene sets (Figures S1A-S1C).

**Figure 1.**
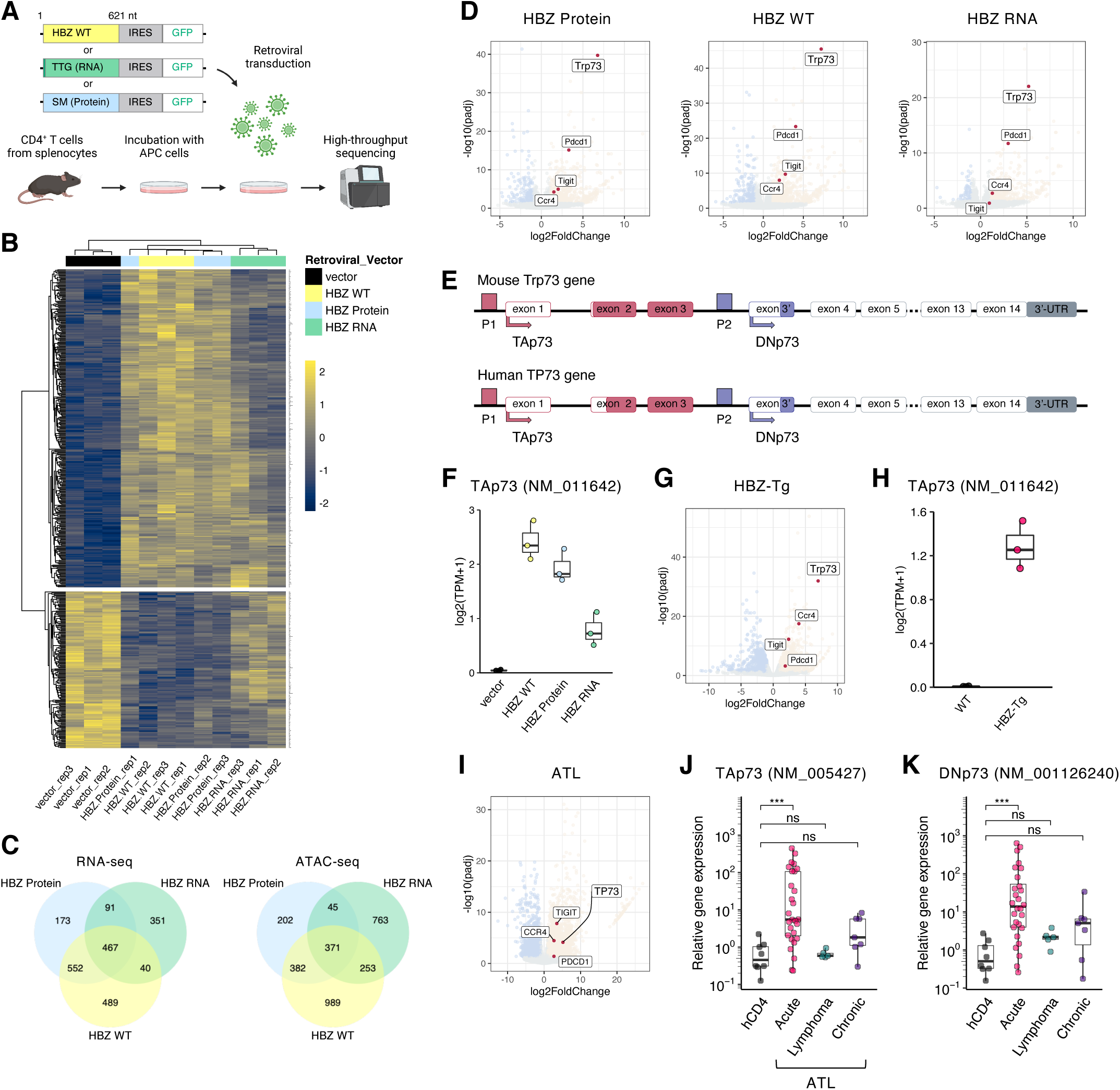
HBZ protein and RNA both upregulate TP73. (A) Schematic diagram for retroviral transfer of HBZ into primary mouse CD4^+^ T cells. Each construct encodes wild-type (WT) HBZ, HBZ RNA (ATG is converted to TTG) or HBZ protein (SM, silent mutations) (Satou *et al*., 2006). (B) A heatmap of the top 500 differentially expressed genes in HBZ WT and its mutants compared to the vector, calculated from RNA sequencing (RNA-seq) data. (C) Shared transcriptomes (left; the number of genes) and open chromatin regions (right) associated with HBZ WT and its mutants are shown in Venn diagrams. (D) Volcano plots of differentially expressed genes. Fold change and adjusted p-value (padj) are plotted for genes that are up-reulgated (yellow) or down-regulated (blue) compared to the vector. (E) TP73 gene maps (upper, mouse; lower, human) depicting the major two isoforms, TAp73 and DNp73, and their promoters. (F) Transcripts per million (TPM) of TAp73 in the transduced cells (n=3). (G) A volcano plot resulting from RNA-seq that compares the CD4^+^ T cells of HBZ-transgenic mice (HBZ-Tg) with those of WT mice (n=3). (H) TPM of TAp73 in HBZ-Tg and WT mouse CD4^+^ T cells (n=3). (I-K) TP73 expression in adult T-cell leukemia-lymphoma (ATL). A volcano plot resulting from RNA-seq that compares CD4^+^ T cells of ATL patients (n=7) with those of healthy donors (n=10) (I). mRNA expression of TAp73 (J) and DNp73 (K) by RT-qPCR in CD4^+^ T cells of ATL patients (acute type, n=32; lymphoma type, n=7; chronic type, n=7) and healthy donors (hCD4; n=6). Results are plotted as mean ± SD, using one-way ANOVA followed by post-hoc Steel test (J and K). ***p < 0.001; ns, not significant. See also Figure S1.

Among the common target genes, we focused on *Trp73* because it was highly expressed in cells containing any of the three *HBZ* constructs (Figure 1D). Because it has alternative promoters, *Trp73* encodes two major isoforms: i) TAp73, a full-length protein with a transactivation (TA) domains and ii) DNp73,an N-terminal truncated protein that lacks the TA domains (Figure 1E) (Melino et al., 2002). Among the two isoforms, *TAp73* was upregulated by *HBZ* (Figure 1F), whereas *DNp73* was not (at least in moue cells, see below) (Figure S1D). Similarly, CD4^+^ T cells of HBZ-Tg mice showed upregulation of *TAp73* but not *DNp73* (Figures 1G, 1H and S1E). We next evaluated clinical samples from patients with ATL, and we found that *TAp73* was upregulated in ATL cells -- and *DNp73* was also upregulated in human ATL cases. Moreover, the upregulation of both transcripts was associated with acute-type ATL, which is the most aggressive clinical subtype (Figures 1I-1K). Altogether, these our observations suggest that the *TP73* gene, particularly *TAp73*, is a target of both HBZ protein and *HBZ* RNA.

TAp73 has been thought to exert cancer-suppressive functions due to its pro-apoptotic effect and genomic integrity maintenance (Stiewe and Pützer, 2000; Tomasini et al., 2008). It thus seemed odd that TAp73 would be upregulated in ATL cells. We therefore analyzed The Cancer Genome Atlas (TCGA) RNA-seq datasets to assess transcriptional profiles of the *TP73* gene in a variety of cancer cells. Surprisingly, the vast majority of registered cancer cells have enhanced expression of TP73, and in particular they have a greater enhancement of expression of *TAp73* than of *DNp73* (Figures S1F-S1H). Thus, we decided to explore any potential oncogenic effects of TAp73 in ATL cells.

### HBZ protein binds to EZH2 and reduces its recruitment to the TAp73 promoter without inhibiting its methyltransferase activity

To investigate the mechanisms for *TAp73* induction by HBZ protein and RNA, we next analyzed the genetic elements in the promoter of *TAp73* using the datasets of chromatin immunoprecipitation sequencing (ChIP-seq) provided by the Encyclopedia of DNA Elements (ENCODE) (Davis et al., 2018; ENCODE.Project.Consortium, 2012). Around the *TAp73* promoter (P1 in Figure 1E), the target regions of two components of PRC2 (EZH2 and SUZ12) were significantly enriched (Figure S2A). Our RNA-seq results also showed that *HBZ* WT and protein, but not RNA, enhanced the same gene sets (PRC2_EZH2_UP.V1_UP) that are upregulated when *EZH2* is knockdown (Figures 2A and S1A), suggesting that HBZ protein might be involved in epigenetic regulation through EZH2. Indeed, immunoprecipitation assays showed that HBZ protein bound directly to EZH2 (Figures 2B and S2B). Experiments using deletion mutants of EZH2 (Figure S2C) revealed that HBZ protein binds to the regions of EZH2 where EZH2 binds to other components of PRC2, but not to the catalytic region including the SET domain (Figure S2D). HBZ protein also co-immunoprecipitated with SUZ12 and EED (Figure S2E). Thus, our findings imply that HBZ protein binds to the PRC2 complex and affects its function.

**Figure 2.**
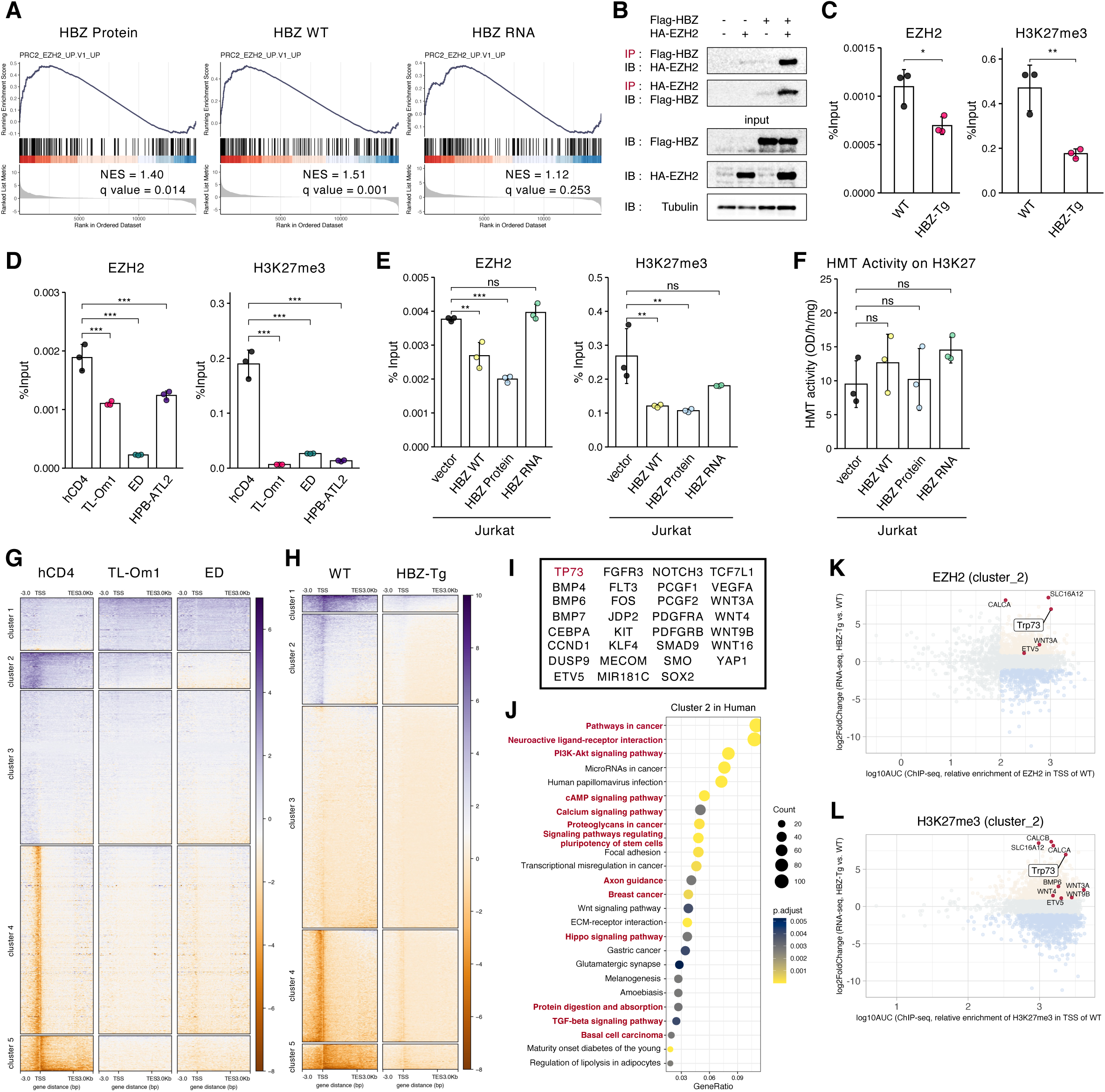
HBZ protein alters EZH2 genome-wide distribution and decreases its binding to the TAp73 promoter. (A) GSEA plots for mouse CD4^+^ T cells transduced with WT or mutant HBZ compared to the vector. The normalized enrichment score (NES) and false discovery rate (FDR) q-value are listed. (B) Immunoprecipitation (IP) with anti-Flag antibody (Flag-HBZ) showing interaction between HBZ protein and EZH2 in 293T cells. IP was analyzed by SDS-PAGE and immunoblotting (IB). (C-E) ChIP qPCR for EZH2 and H3K27me3 in the TAp73 promoter region. The %Input is shown for WT or HBZ-Tg mouse CD4^+^ T cells (C), human CD4^+^ T cells (hCD4) or ATL cell lines (D), and Jurkat cells with stable transduction of WT or mutant HBZ (E) (n=3). (F) Histone methyltransferase (HMT) activity on H3K27 among transduced Jurkat cells (n=3). (G and H) Heatmaps for genomic regions with enriched ChIP-seq scores for H3K27me3. The score for each DNA region was calculated based on ATL cell lines (TL-Om1 and ED) relative to healthy human donor CD4^+^ T cells (hCD4) (G) or HBZ-Tg mouse CD4^+^ T cells relative to WT (H). (I) Representative genes found in cluster 2 that were shared between the results from human and mouse cells in G and H respectively. (J) Results of KEGG pathway analysis using the genes in cluster 2 for humans. Statistical values and gene counts calculated by the clusterProfiler are shown. (K and L) Scatter plots of mouse cluster 2 genes resulting from combinational analysis of RNA-seq (HBZ-Tg mouse CD4^+^ T cells relative to WT) and ChIP-seq (relative enrichments in WT) for EZH2 (K) and H3K27me3 (L). Results are plotted as mean ± SD, using Student’s t test (C) or one-way ANOVA with post-hoc Dunnet test (D–F). *p < 0.05, **p < 0.01, ***p < 0.001; ns, not significant. See also Figure S2; Tables S1 and S2.

The epigenetic status of *TAp73* promoter regions in ATL cells and in CD4^+^ T cells of HBZ-Tg mice was evaluated with ChIP quantitative real-time PCR (ChIP-qPCR) and ChIP-seq. Of note, EZH2 recruitment and H3K27me3 were reduced, and H3K27 acetylation (H3K27ac) was increased in the *TAp73* promoter in both HBZ-Tg mouse CD4^+^ T cells and ATL cell lines compared to their corresponding controls (Figures 2C, 2D and S2F). This reduction in EZH2 recruitment would be expected to result in increased transcription of *TAp73*, which would explain why *TAp73* expression levels are elevated in HBZ-containing cells. The idea that reduced levels of EZH2 and H3K27me3 in the *TAp73* promoter were caused by HBZ protein, but not RNA, was confirmed using HBZ-transduced Jurkat cells (Figure 2E). We also found that *HBZ* WT and its mutants did not affect EZH2 enzymic activity (Figure 2F). All these observations are consistent with the hypothesis that HBZ protein binds to the PRC2 complex but not to the catalytic domain of EZH2.

### HBZ protein dysregulates genome-wide distribution of H3K27me3

Our findings indicate that HBZ protein changes the transcriptional profiles of some genes by binding to EZH2 without affecting its enzymatic activity. Therefore, we analyzed the genome-wide distribution of EZH2 and H3K27me3 in ATL cells and HBZ-expressing cells. As expected, the accumulation of H3K27me3 in transcription start sites (TSSs) changed in a wide variety of genes in both ATL cell lines and CD4^+^ T cells of HBZ-Tg mice (Figures 2G, 2H, and Tables S1 and S2). Multiple genes were found to be transcriptionally repressed due to accumulation of the trimethylation (cluster 1 in Figures 2G and 2H), including tumor suppressor genes such as *NDRG2*, in line with previous reports (Fujikawa *et al*., 2016; Nakahata et al., 2014). Concurrently, there were numerous genes in which H3K27me3 in TSS’s was decreased, indicating that they were transcriptionally activated (cluster 2 in Figures 2G and 2H). Of note, many genes in cluster 2, including *TAp73*, were shared by both ATL cells and the CD4^+^ T cells of HBZ-Tg mice (Figure 2I), and pathway analysis indicated that they are associated with various oncogenic pathways (Figures 2J and S2G). *TAp73* upregulation in HBZ-Tg CD4^+^ T cells was significantly correlated with reduced levels of EZH2 and H3K27me3 in its promoter (Figures 2K and 2L), supporting the idea that EZH2 is a significant regulator of *TAp73* expression.

### HBZ RNA-enhanced BATF3 elicits activation of both the TAp73 and DNp73 promoters in human cells

Thus far we have discussed the epigenetic upregulation of *TAp73* transcription by HBZ protein via its effects on EZH2. *HBZ* RNA is thought to upregulate *TAp73* by a different mechanism. We found that an open chromatin region was present about 3kb upstream from the *TAp73* TSS, especially in *HBZ* RNA and WT-transduced cells (highlighted by gray; Figure S3A). H3K27ac ChIP-seq of HBZ-Tg CD4^+^ T cells revealed that transcription was activated in this region, where BATF3 and IRF4 can be recruited (Figure S3A, right). Corroborating these findings, *BATF3* transcription was drastically upregulated by *HBZ* RNA, compared with *HBZ* WT and protein (Figure 3A) as previously reported (Nakagawa et al., 2018). Additionally, it was found that the target sequences of BATF are also activated by *HBZ* RNA (Figure 3B). We identified a similar region within the *TP73* gene body in the human genome (highlighted by gray; Figure 3C). Recruitment of both BATF3 and IRF4 to this region was confirmed by ChIP-seq and ChIP-qPCR (Figures 3C, S3B and S3C) (Nakagawa *et al*., 2018). BATF3 is known to alter the expression of ATL-associated genes in concert with IRF4 (Ishio et al., 2022; Nakagawa *et al*., 2018). Thus, we next performed promoter assays using the identified region upstream of human *TAp73*. As expected, *HBZ* RNA induced this promoter activity (Figure 3D). The position of the target motifs of BATF3 and IRF4 is closer to both the *TAp73* and *DNp73* promoters in the human genome (Figure 3C, left) than in the mouse genome (Figure S3A, left). Intriguingly, the promoters of both human *TAp73* and *DNp73* were activated by BATF3 and IRF4 (Figures 3E, 3F, and S3D, S3E), suggesting that *HBZ* RNA induces *DNp73* in human cells, even though it cannot in mouse cells (Figure S1D). Taken together, our data show that HBZ protein and RNA induce *TAp73* in different manners, and *HBZ* RNA activates *DNp73* in human cells but not in mouse cells. These findings could collectively explain why both *TAp73* and *DNp73* were upregulated in patients with acute ATL (Figures 1J and 1K).

**Figure 3.**
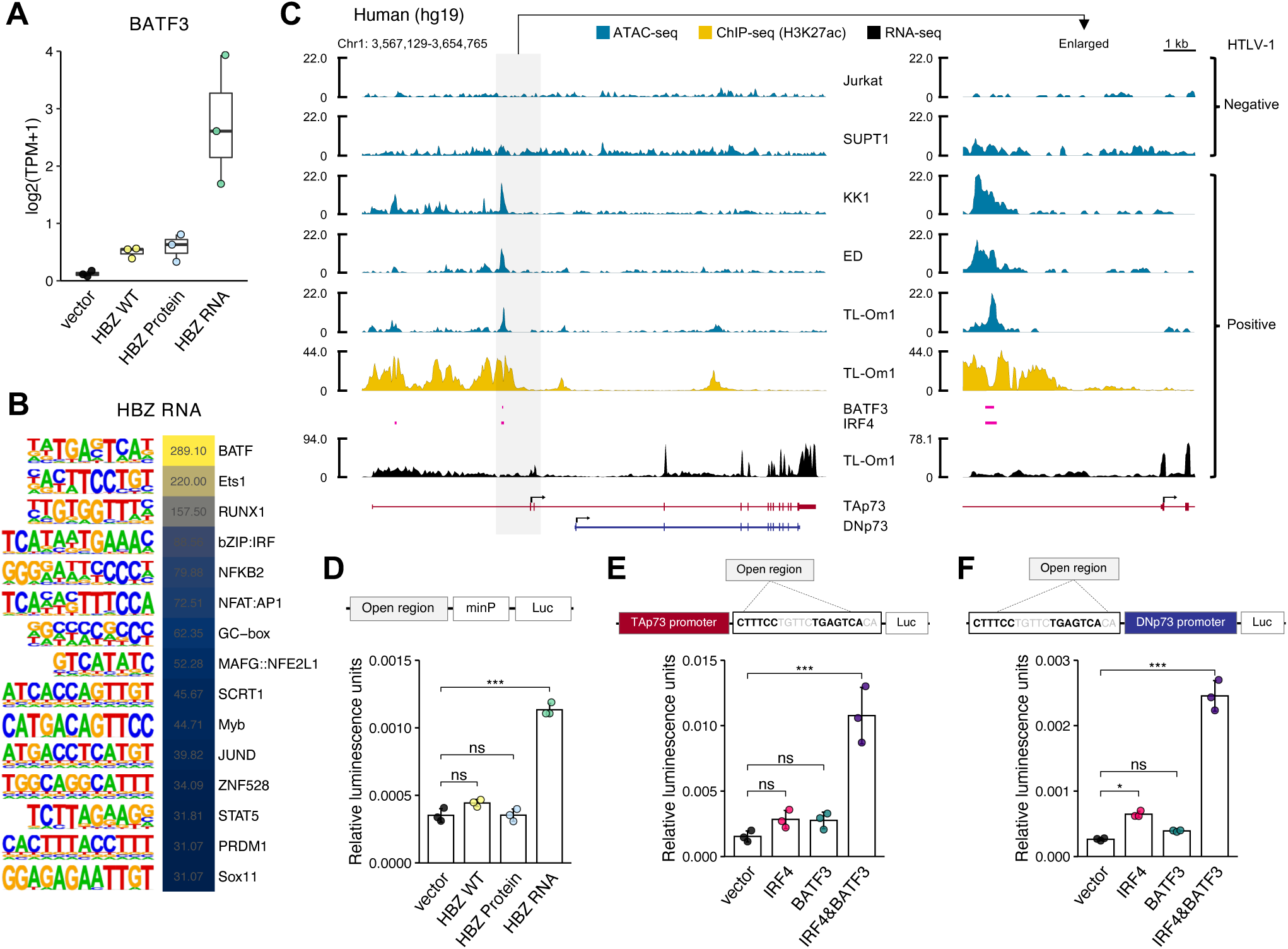
HBZ RNA enhances both the TAp73 and DNp73 promoters via BATF3-IRF4 transcriptional machinery. (A) TPM of BATF3 in HBZ-transduced murine CD4^+^ T cells (n=3). (B) The top-ranked enriched motifs from (A) with their log_2_ p-values from the findMotifsGenome results (HOMER). (C) Chromatin accessibility (ATAC-seq), H3K27ac enrichment, BATF3/IRF4 binding regions (ChIP-seq; SRX2548278 and SRX2548284) (Nakagawa *et al*., 2018) and transcripts (RNA-seq) of the TP73 gene in HTLV-1 negative or positive human T-cell lines. (D) Promoter assays of the HTLV-1-specific open region identified in Figure 3C (hg19 genome region of chr1:3593076-3594185) with a minimal promoter (minP) in 293 cells co-transfected with WT or mutant HBZ. The open region was inserted into pNL3.2.CMV after cloning of the genome region as shown in a schematic of construct. (E-F) The IRF4/AP-1 motifs identified within the open region were subjected to promoter assays with IRF4 and/or BATF3 induction: for the promoter of TAp73 (E) and the promoter of DNp73 (F) (n=3). A schematic of the assay construct is shown above the corresponding bar plot. Results are plotted with mean ± SD, using one-way ANOVA with post-hoc Dunnet test (D–F). *p < 0.05, ***p < 0.001; ns, not significant. See also Figure S3.

### TAp73 activates EZH2 transcription

Thus far we have shown that *HBZ* RNA appears to increase BATF-activated *TAp73* transcription, while HBZ protein activates *TAp73* by interacting with EZH2. Interestingly, *HBZ* also appears to cause elevated levels of EZH2 -- by a mechanism we can now explain. High-level expression of EZH2 with accumulation of H3K27me3 has been previously reported in ATL cells (Fujikawa *et al*., 2016; Sasaki *et al*., 2011; Yamagishi *et al*., 2012). *HBZ* and *tax*, another HTLV-1 viral oncogene, have been thought to affect EZH2 regulation (Akkouche et al., 2021; Yamagishi et al., 2018), but the details have been unclear. Surprisingly, we found that TAp73 was recruited to the promoter region of the *EZH2* gene in ATL cell lines (Figures 4A and 4B). Luciferase assays revealed that TAp73, but no other TP73 isoform, enhances *EZH2* promoter activity (Figure 4C). Indeed, ATL cells endogenously express both TAp73 and EZH2 proteins (Figure 4D). As previously reported (Yamagishi et al., 2019), we confirmed the upregulation of *EZH2* gene in ATL patients, while finding no significant difference in *EZH1* expression (Figure 4E). Importantly, a positive correlation between *TAp73* and *EZH2*, but not *EZH1*, was observed in ATL patients (Figure 4F) and patients with various cancers in TCGA datasets (Figure 4G), suggesting that TAp73 is associated with the regulation of *EZH2* gene expression not only in ATL but also in other types of cancer.

**Figure 4.**
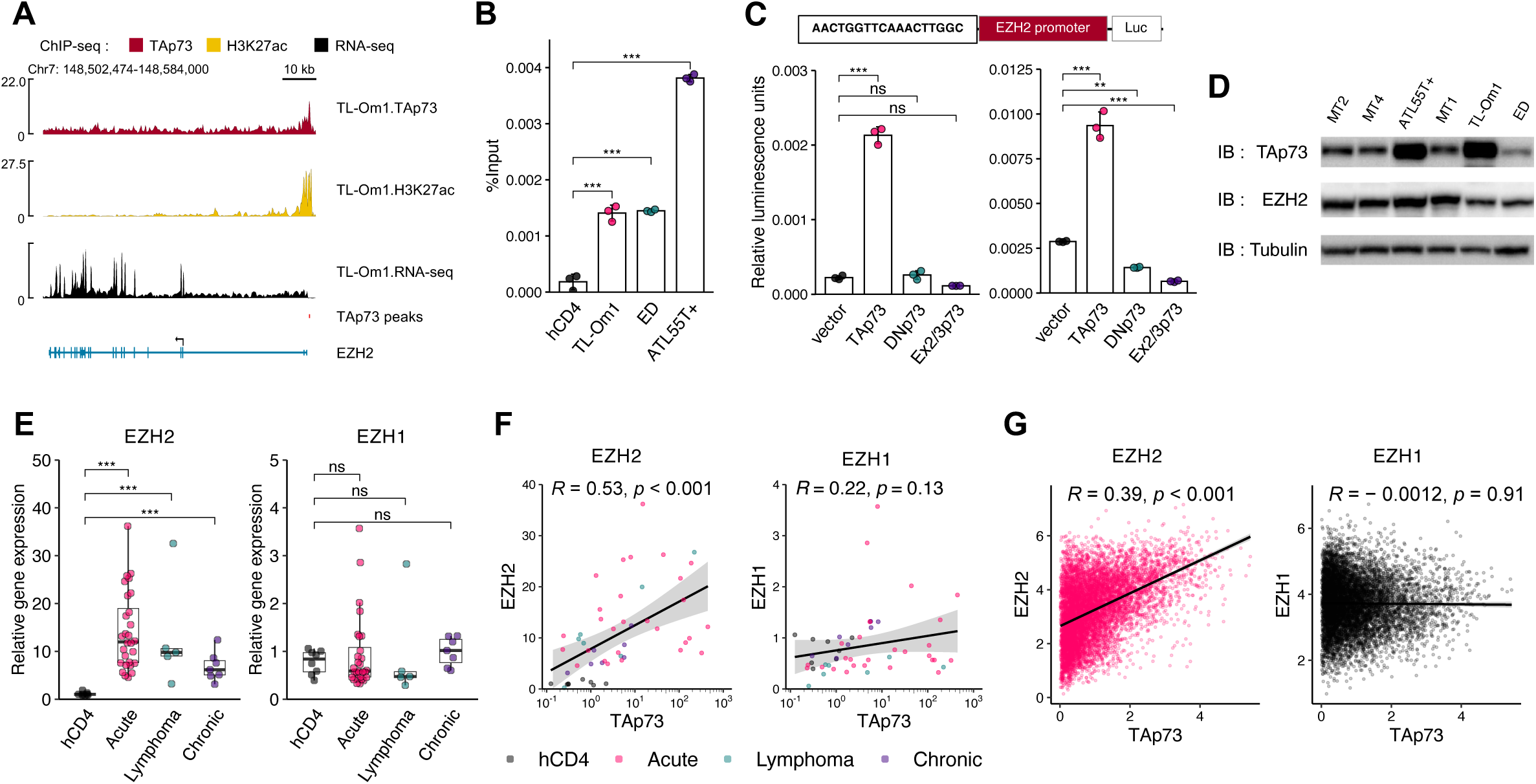
TAp73 transcriptionally induces EZH2 gene expression. (A) TAp73 and H3K27ac enrichment (ChIP-seq) and transcripts (RNA-seq) of the EZH2 gene in TL-Om1 cells. (B) ChIP-qPCR of ATL cell lines relative to healthy human donor CD4^+^ T cells (hCD4) (n=3). (C) Promoter assays using the TP73 motif identified within TAp73 peaks by the ChIP-seq experiment shown in (A). Relative luciferase activities with the expression of various TP73 isoforms in 293 (left) and Jurkat (right) cells (n=3). A schematic of the assay construct is shown above the bar plots. (D) Immunoblots of TAp73, EZH2 and Tubulin in HTLV-1-infected cell lines. (E) mRNA expression of EZH2 (left) and EZH1 (right) by RT-qPCR in hCD4 (n=6) and ATL cells (acute type, n=32; lymphoma type, n=7; chronic type, n=7). (F and G) Correlation between TAp73 expression and expression of EZH2 (left) or EZH1 (right) in hCD4 and ATL cells (F) and TCGA data (G). Results of Pearson correlation analysis are plotted. Results are plotted as mean ± SD, using one-way ANOVA with post-hoc Dunnet (B and C) or Steel test (E). **p < 0.01, ***p < 0.001; ns, not significant.

### Inactivation of TAp73 induces cell death and acidification by lactate accumulation in ATL cells

To evaluate the significance of our findings in *HBZ*-mediated pathogenesis in vivo, we generated two strains of *Trp73* knockout (KO) mice, *TAp73*−/− and *DNp73*−/− (Figures 5A and S4A), and we crossed them with HBZ-Tg mice. HBZ-Tg mice develop systemic inflammation, such as dermatitis and bronchitis (Satou *et al*., 2011). Compared to the frequency of dermatitis in HBZ-Tg mice, the frequency of dermatitis was significantly reduced in HBZ-Tg/*TAp73*−/− mice, but not in HBZ-Tg/*DNp73*−/− mice (Figure 5B). These findings support the interpretation that in mice, the functions of TAp73, not DNp73, are of primary importance in the pathogenicity of *HBZ*.

**Figure 5.**
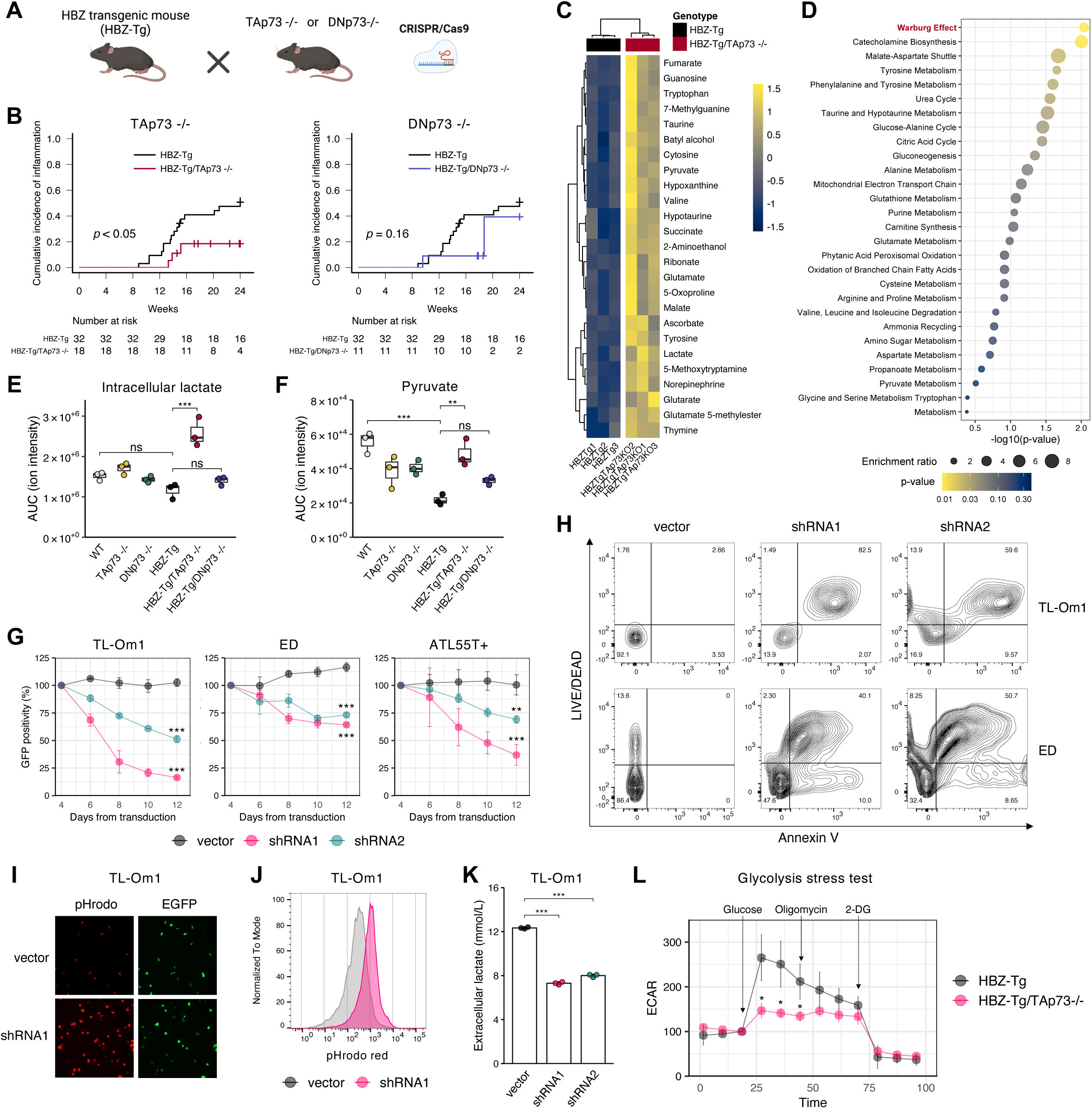
TAp73 inactivation causes ATL cell death and intracellular acidification due to lactate accumulation. (A) Mating diagram of HBZ-Tg mice with TP73 knockout mice (TAp73−/− or DNp73−/−) created by CRISPR/Cas9. (B) Cumulative incidence of skin inflammation in HBZ-Tg mice (n=32) compared to HBZ-Tg/TAp73−/− mice (left; n=18) or HBZ-Tg/DNp73−/− mice (right; n=11). The p values determined by Gray test are shown. (C and D) Changes in metabolites between HBZ-Tg and HBZ-Tg/TAp73−/− mouse CD4^+^ T cells. Shown are the top 25 differentially altered metabolites (C) and enriched metabolite sets (D). (E and F) Intracellular lactate (E) and pyruvate (F) levels in the murine CD4^+^ T cells (n=3). Calculated ion intensities (area under the curve: AUC) are shown. (F) GFP competition assay of ATL cell lines with TP73 knockdown (n=3). Lentiviral vectors for knockdown encoded EGFP. The date of lentiviral transduction was counted as day 1. Values in comparison to the vector are shown. (G) Cell viability and apoptosis assay with TP73 knockdown. On day 8 after transduction, ATL cells were analyzed by flow cytometry. Representative dot plots are shown. (I and J) Intracellular pH assessed by pHrodo Red AM in TL-Om1 cells on day 8 after transduction. Fluorescence microscopy photographs with EGFP (indicates lentivirus-transduced cells) and pHrodo Red AM (I) and flow cytometry results (J). (K) Extracellular lactate in TL-Om1 cell cultures on day 8 after transduction (n=3). (L) Metabolic flux assay of CD4^+^ T cells from HBZ-Tg or HBZ-Tg/TAp73−/− mice (n=3). Glucose, oligomycin and 2-deoxyglucose (2-DG) were injected as the indicated time points. Results are plotted as mean ± SD, using the Student’s t test (L), one-way ANOVA with post-hoc Turkey (E) or Dunnet test (G and K). *p < 0.05, **p < 0.01, ***p < 0.001; ns, not significant. See also Figure S4.

TAp73 was previously shown to enhance both glycolysis and the pentose phosphate pathway (PPP) by activating transcription of the gene encoding the enzyme that catalyzes each pathway’s rate-limiting step (Du et al., 2013; Li et al., 2018). Indeed, HBZ-Tg mice showed higher expression levels of *G6pdx* and *Pfkl* (Figure S4B). Our HBZ-Tg/*TAp73*−/− mouse models are useful for assessing the significance of TAp73 in glucose metabolism and its involvement in *HBZ*-mediated pathogenesis *in vivo*. Metabolomics analysis showed that CD4^+^ T cells from HBZ-Tg vs. HBZ-Tg/*TAp73*−/− mice had markedly different metabolite profiles (Figure 5C). HBZ-Tg/*TAp73*−/− cells showed similar profiles to WT ones (Figure S4C), suggesting that *TAp73* KO cancels *HBZ*-driven metabolic alteration. When we analyzed the metabolites altered differentially between HBZ-Tg and HBZ-Tg/*TAp73*−/− mice, a metabolite set related to the Warburg effect was highly enriched (Figure 5D). Unexpectedly however, we found that the intracellular metabolites of glycolysis, such as lactate, pyruvate, and G6P, were higher in HBZ-Tg/*TAp73*−/− mice than HBZ-Tg mice (Figures 5E, 5F, and S4D). This was an unexpected result, because if glycolysis and/or PPP are suppressed, then their end products, would be reduced as well.

To examine these findings in more detail, we next performed knockdowns in ATL cell lines using two short hairpin RNAs: shRNA1 targets only *TAp73*, while shRNA2 works against all isoforms of *TP73* (Figures S4E–S4G). As expected, we found that *EZH2* mRNA was reduced by knockdown of *TAp73* (Figure S4H). Both shRNAs induced necrotic (or late apoptotic) cell death, and intriguingly, this effect was especially robust for shRNA1 (Figures 5G and 5H). Intracellular pH became more acidic (Figures 5I, 5J, and S4I, S4J), and the concentration of extracellular lactate was reduced (Figures 5K and S4K). We then performed a metabolic flux assay on the glycolytic system. TAp73 KO in HBZ-Tg mice led to a markedly reduced extracellular acidification rate (ECAR; Figure 5L). Taken together, these observations indicate that inactivation of *TAp73* suppresses lactate excretion out of ATL cells, resulting in its intracellular accumulation and cell death.

### Lactate transporters induced by TAp73 are novel therapeutic targets for ATL

To test the hypothesis that TAp73 regulates lactate transport, we re-analyzed the results of ChIP-seq for TAp73. In ATL cell lines, TAp73 bound to the promoter regions of *SLC16A1* and *SLC16A3*, which encode the lactate transporters MCT1 and MCT4 respectively (Figures 6A and 6B). TP73-binding motifs were identified in the extracted ChIP-seq peaks in both genes (red bars in Figure 6A). A promoter assay (Figure S5A) revealed that TAp73, but not other isoforms of TP73, enhanced the promoter activities of both the *SLC16A1* and *SLC16A3* genes (Figures 6C and S5B). Corroborating these results, *SLC16A1* and *SLC16A3* were both upregulated in ATL patients (Figure 6D), whereas the other MCT family genes, *SLC16A7* (encoding MCT2) and *SLC16A8* (encoding MCT3) were not (Figure S5C). Furthermore, both *SLC16A1* and *SLC16A3* expression were positively correlated with *TAp73* expression in ATL cells (Figure 6E). As expected, *TAp73* knockdown in ATL cells resulted in decreased expression of both the *SLC16A1* and *SLC16A3* genes (Figures S5D and S5E). TCGA datasets also demonstrated that *SLC16A1* and *SLC16A3* (but not *SLC16A7* and *SLC16A8*) were dominantly upregulated in multiple cancer types (Figures S5F-S5I). Positive correlations between the expression levels of *TAp73* and *SLC16A1* or *SLC16A3* were also found in those cancers (Figure 6F). Notably, the signatures-high patients had a poorer prognosis with statistical significance across acute myeloid leukemia and multiple solid cancers (Figure 6G). These observations imply that TAp73-MCT1/4 metabolic regulation is related to the prognosis of various cancers.

**Figure 6.**
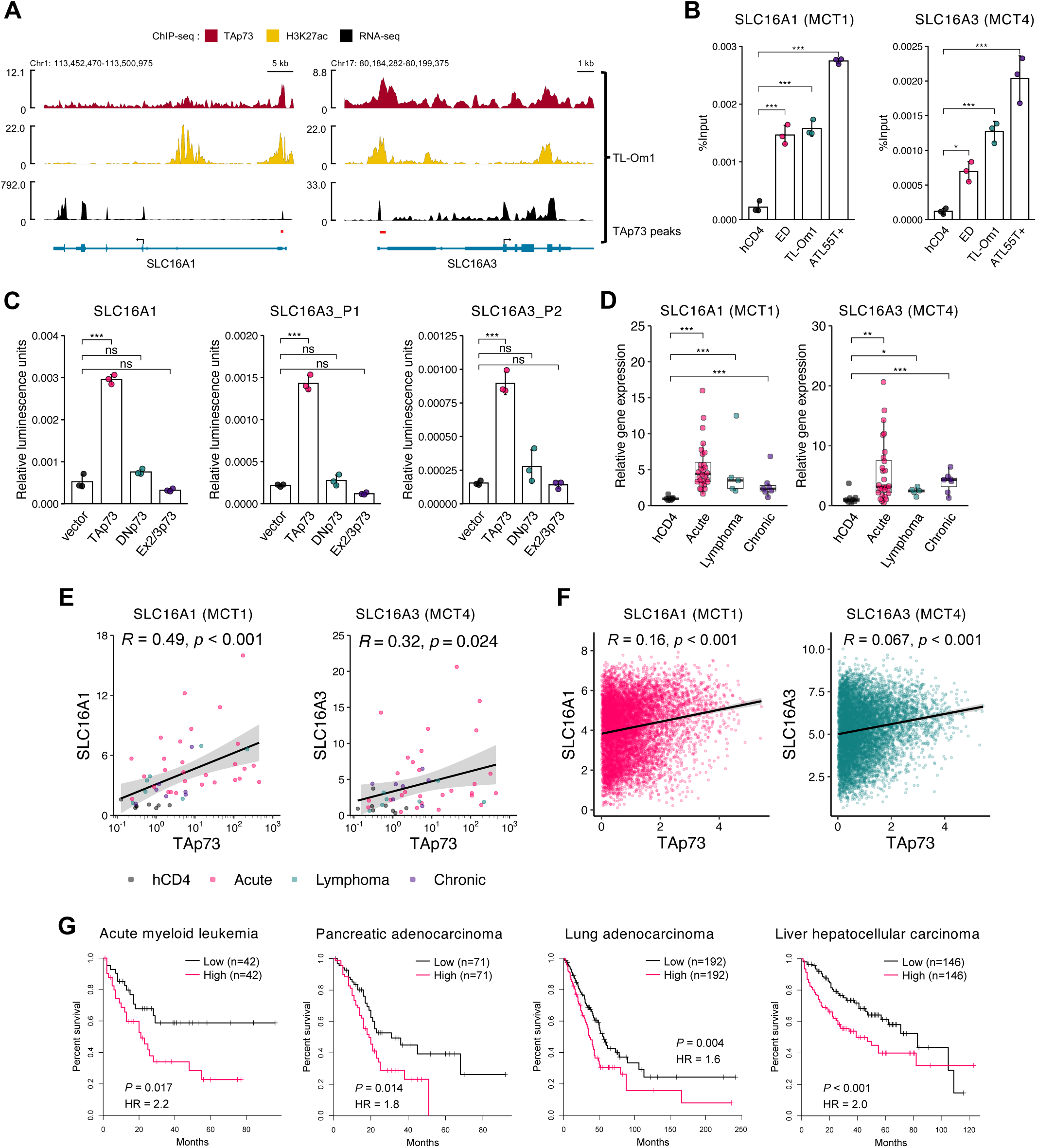
TAp73 upregulates both the SLC16A1 and SLC16A3 genes that encode lactate transporters. (A) TAp73 and H3K27ac enrichments (ChIP-seq) and transcripts (RNA-seq) of the SLC16A1 (left) and SLC16A3 (right) genes in TL-Om1 cells. (B) ChIP qPCR for the SLC16A1 (left) and SLC16A3 (right) promoters in ATL cell lines and human CD4^+^ T cells (hCD4) (n=3). (C) Promoter assays using theTP73 motif identified within the TAp73 peaks shown in (A). Relative luciferase activities for the SLC16A1 promoter (left) and the SLC16A3 promoter (center and right) were examined with TP73 isoform expression in 293 cells. (n=3). Schematics of the assay plasmids are shown in Figure S5A. (D) mRNA expression of SLC16A1 (left) and SLC16A3 (right) measured by RT-qPCR in hCD4 cells (n=6) and ATL patients (acute type, n=32; lymphoma type, n=7; chronic type, n=7). (E and F) Correlation between TAp73 expression and expression of SLC16A1 (left) or SLC16A3 (right) in hCD4 cells and ATL patients (E) and TCGA data (F). (G) Overall survival of patients with TAp73, SLC16A1 and SLC16A3 high or low based on TCGA data (RNA-seq). Statistical values are shown with hazard ratio (HR). Results are plotted with mean ± SD, using one-way ANOVA with post-hoc Dunnet (B, C) or Steel test (D). *p < 0.05, **p < 0.01, ***p < 0.001; ns, not significant. See also Figure S5.

We next evaluated the therapeutic efficacy of syrosingopine, which is an MCT1/4 dual inhibitor (Benjamin et al., 2016; Benjamin et al., 2018) on ATL cells, and found that it induced cell death in a dose-dependent manner (Figures 7A-7C, and S6A, S6B). Importantly, syrosingopine treatments led to intracellular accumulation of lactate (Figure 7D) and reduced the concentration of extracellular lactate (Figure 7E) in ATL cells. Syrosingopine significantly suppressed the growth of ATL cells transplanted into immunodeficient NOD/SCID/IL2Rg^null^ (NSG) mice (Figures 7F-7H). Taken together, these results show that MCT1/4 dual inhibition yielded antitumor effects by blocking lactate excretion from ATL cells *in vitro* and *in vivo*.

**Figure 7.**
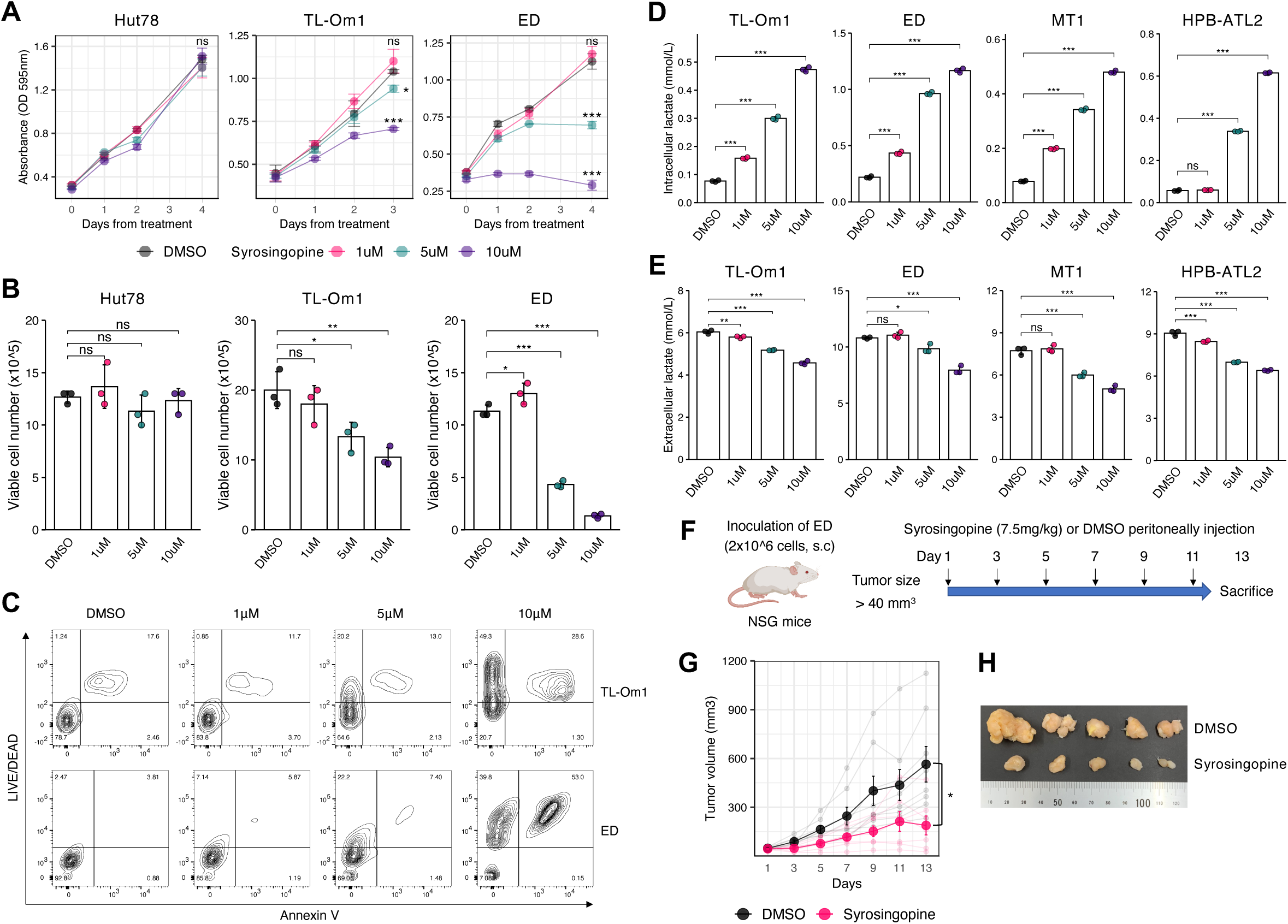
Efficacy of the MCT1/4 inhibitor syrosingopine on ATL cells. (A-C) Cell proliferation assay (A), viable cell numbers (B), and flow cytometric analysis of apoptosis and dead cells at day 4 (C; representative dot plots) for HTLV-1-unifected T cells (Hut78) or ATL cells (TL-Om1 and ED) treated with syrosingopine (1μM, 5μM and 10μM; n=3). Values in comparison to the DMSO group are shown. (D and E) ATL cells expel less lactate when treated with syrosingopine. Lactate was examined intracellularly (D) or extracellularly (E) after 4 days of syrosingopine treatment. (F-H) Therapeutic efficacy of syrosingopine in NSG mice after inoculation with ATL cells. Experimental outline for the *in vivo* experiment to compare syrosingopine with DMSO (F). Tumor volumes were measured every 2 days in NSG mice treated with syrosingopine (n=7) or DMSO (n=8) (G). Representative tumors resected from the mice were shown in (H). Results are plotted as mean ± SD, using one-way ANOVA with post-hoc Dunnet test (A, B, D and E) or Student’s t test (G). *p < 0.05, **p < 0.01, ***p < 0.001; ns, not significant. See also Figure S6.

## Discussion

In many cancer tissues, the metabolism of glucose is often dysregulated in order to fuel cell proliferation. Activation of several protooncogenes and mutations of certain tumor suppressor genes have been shown to reprogram the glycolysis process (DeBerardinis et al., 2008; Jones and Thompson, 2009). Myc increases expression of many metabolic enzymes, including LDH-A, and several enzymes required for nucleotide biosynthesis (Jones and Thompson, 2009). It has been also reported that constitutive activation of Akt increases the surface expression of glucose transporters and stimulates expression and enzyme activity of the factors associated with glycolysis (DeBerardinis *et al*., 2008). Multiple lines of evidence indicate that ATL shares multiple hallmark capabilities with other cancers (Ma et al., 2016; Mesri et al., 2014); however, there are few studies reporting the involvement of altered glucose metabolism in the oncogenic activity of HTLV-1. In this study, we show that an isoform of the *TP73* gene containing the transactivation domain, *TAp73*, is induced by *HBZ* and plays critical roles in the excretion of lactate, which is a waste metabolite of glycolysis (Rabinowitz and Enerbäck, 2020). Several studies previously showed that TAp73 transactivates genes associated with glycolysis, such as *GP6D* and *PFKL*, suggesting that TAp73 might have a role in the Warburg effect (Li *et al*., 2018). Expression of these genes was also upregulated in *HBZ* transgenic cells (Figure S4B), and more notably, the expression of the lactate transporters MCT1 and MCT4 was significantly upregulated, suggesting that TAp73 both accelerates the glycolytic cascade and boosts the efflux of metabolites out of the cells. Interestingly, the positive correlation between *TAp73* and *SLC16A1/3* expression is observed in other cancer types, and this signature is related to their prognosis with statistical significance (Figures 6F and 6G). These observations imply that promotion of lactate excretion by TAp73-iduced MCT1/4 is a common feature of many cancers.

We also found that in addition to inducing glycolysis and lactate transport, TAp73 induces *EZH2* transcription (Figures 4A-4C). Previous studies have shown that EZH2 is highly expressed in ATL cells and alters their epigenetic landscape to repress the expression of several tumor suppressor genes such as *NDRG2* and *ZEB1* (Fujikawa *et al*., 2016). Since inhibitors of EZH2 have been proven to be effective against ATL cells (Yamagishi *et al*., 2019), EZH2 is recognized as one of the key molecules involved in ATL leukemogenesis. Our results suggest that transcriptional upregulation of *TAp73* by *HBZ* is an important mechanism for inducing constitutive expression of *EZH2* in ATL cells and even in HTLV-1-infected cells. Surprisingly, we also identified a strong correlation between *TAp73* and *EZH2* transcription levels in the TCGA dataset (Figure 4G), suggesting that TAp73 is a common transactivator of *EZH2* in many types of cancer. Thus TAp73 is associated with the “nonmutational epigenetic reprogramming” described in the recently revised Hallmarks of Cancer (Hanahan, 2022).

Clonal expansion of infected cells is crucial for the persistent infection of HTLV-1 in vivo, and eventually leads to malignant transformation of infected cells. *HBZ* plays important roles in the proliferation and survival not only of ATL cells but also of infected nonleukemic cells. We and others have reported the distinct functions of HBZ protein and *HBZ* RNA (Gazon et al., 2020; Mitobe *et al*., 2015; Satou *et al*., 2006). RNAs imbued with both coding and non-coding functions are known to exist in bacteria, plants, and animals and are referred to as “bifunctional RNAs” or “coding and non-coding RNAs (cncRNAs)” (Sampath and Ephrussi, 2016). In mammals, several mRNAs have been proven to harbor noncoding functions independent of the proteins they encode. For example, *Steroid receptor activator* (*SRA*), a functional RNA, and SRAP, a translated protein, counteract each other in myogenic differentiation (Hubé et al., 2010). A well-known tumor suppressor gene, *TP53*, also has coding and non-coding functions; *p53* mRNA inactivates the E3 ubiquitin ligase MDM2, a negative regulator of p53 protein, and consequently controls p53 responses under genotoxic stresses (Candeias et al., 2008). Each case of bifunctional RNA is thought to be a native machinery for fine-tuning their own activities. In contrast, *HBZ* RNA and protein regulate the function and/or expression of numerous host factors and promote the clonal proliferation of HTLV-1-infected cells. Here, we show that both HBZ protein and *HBZ* RNA strongly induce *TAp73* expression through different mechanisms: HBZ protein diminishes recruitment of EZH2 to the *TAp73* promoter, whereas *HBZ* RNA upregulates *BATF3*, which activates the *TAp73* promoter. Thus HTLV-1 has redundant mechanisms to induce TAp73. *HBZ* RNA was found to change its subcellular localization depending on its promoter activity (Ma *et al*., 2021), and it is important for *HBZ* RNA to localize in the nucleus to promote cell growth. Considering that translation into protein is generally executed in the cytoplasm (Buxbaum et al., 2015; Yan et al., 2016), it is possible that *HBZ* RNA and HBZ protein are differentially expressed depending on the activity of the HBZ promoter. Redundant mechanisms for TAp73 induction by *HBZ* RNA and protein might be useful to maintain TAp73 expression regardless of cell status. Recently, it has been reported that the intragenic super-enhancer is present in *TP73* gene in ATL cells, and it contributes to the high expression of *TP73* and cell growth (Ong et al., 2022).

Since HBZ protein reduces the level of H3K27me3 in the same genomic region in human T cells (Figures 2E and S2F), it’s suggested that modulation of the epigenetic status by HBZ is involved in the formation of the *TP73* intragenic super-enhancer, which supports its continuous expression.

The role of TP73 in oncogenesis has been mainly associated with the expression of DNp73, a dominant negative isoform with anti-apoptotic properties (Melino *et al*., 2002). Our expression analysis revealed increased expression of both *TAp73* and *DNp73* in ATL cells (Figures 1J and 1K), but we focused on the unexpected result that knockdown of *TAp73* alone reduced proliferation of the ATL cell lines we used (Figure 5G). Previous genomic studies reported that ~20% of ATL cases lack the transactivation domain of the *TP73* gene due to structural aberration (Kogure et al., 2021), suggesting that in those cases, the anti-apoptotic effect of DNp73 is more important for the survival of ATL cells than the transactivating properties of TAp73.

Our results suggest that *BATF3*, induced by *HBZ* RNA, is associated with the activation of its target motif near to the human *TP73* gene (Figures 3C-3F). Similar AICE motifs are absent near to the mouse *Trp73* gene. This difference is a possible reason for the fact that *DNp73* is induced by *HBZ* RNA in human T cells (Figure 1K) but not in mouse T cells (Figures S1D and S1E). The fact that *HBZ* RNA does not induce *DNp73* expression in mice may explain why *TAp73*, not *DNp73*, was important in the development of dermatitis (Figure 5B). In humans the story may be more complicated. We have shown that TAp73 plays an important role in ATL. DNp73 was also reported to function as an oncogenic protein via its inhibitory effect on TAp73- or TP53- mediated apoptosis (Engelmann et al., 2015; Grob et al., 2001), and we found that DNp73, like TAp73, responds to the induction of *BATF3* by *HBZ* RNA. At this moment, the molecular mechanisms for induction of *BATF3* by *HBZ* RNA remain unknown, and they will need to be elucidated in future.

In summary, TAp73, induced by HBZ, contributes to both the Warburg effect and epigenetic reprogramming in HTLV-1-infected cells via induction of MCT1/4 and EZH2, respectively. These signatures are commonly observed in multiple types of cancer, implying that these molecules are promising targets for wide-spectrum anti-cancer strategies.

## Acknowledgments

We thank Dr. Linda Kingsbury for proof-reading of this manuscript. We also thank Chiho Onishi, Miho Matsumoto, Saiko Tokunaga, Chiaki Ohama and Ryu Yamashita for their technical assistance with the experiments. This work was supported by a grant from the Project for Cancer Research And Therapeutic Evolution (P-CREATE) (20cm0106306h0005 to M. M.), the Research Program on Emerging and Re-emerging Infectious Diseases (20fk0108088h0002 to M. M.) from the Japan Agency for Medical Research and Development (AMED); Science and Technology Platform Program for Advanced Biological Medicine (21am0401003h0003 to J.Y.) from AMED; JSPS KAKENHI (19H03689 to M.M.) and JSPS KAKENHI (20H03514 to J.Y.). This study was also supported in part by the JSPS Core-to-Core Program A, Advanced Research Networks.

## Author contributions

K.Toyoda., J.Y. and M.M. conducted the research design. K.Toyoda., O.H., M.S., M.W. and T.S. performed the experiments. K.Toyoda., Y.A., A.T., O.H., M.W. and D.K. conducted the data analysis. Y.A., K.Tsujita., K.N., K.Y., H.M. and K.O. contributed to discussions. K.Toyoda., J.Y. and M.M. wrote the manuscript. J.Y. and M.M. supervised the study.

## Methods

### RESOURCE AVAILABILITY

#### Lead contact

Further information and requests for materials should be directed to the lead contact: Jun-ichirou Yasunaga.

#### Materials availability

This study did not generate new unique reagents.

#### Data and code availability

Data of RNA-seq, ATAC-seq and ChIP-seq in this study have been deposited at the DDBJ database and are publicly available as of the date of publication. Any additional information required to reanalyze the data reported in this paper is available from the lead contact upon request.

### EXPERIMENTAL MODEL AND SUBJECT DETAILS

#### Clinical samples

Peripheral blood mononuclear cells (PBMCs) were isolated from healthy donors, acute- and chronic-type ATL patients with 3% dextran and the density gradient medium Ficoll-Paque Plus (GE Healthcare). For lymphoma-type ATL, we obtained primary ATL cells from the enlarged lymph nodes for subsequent experiments, since the tumor cells are not detectable in the blood. Isolation of CD4^+^ T cells was carried out by negative selection using the Human CD4 T Lymphocyte Enrichment Set-DM (BD Biosciences). All patient samples were obtained with informed consent for research use. This study was conducted in accordance with the principles in the Declaration of Helsinki, and was approved by the Institutional Ethics Committee of Kumamoto University (approval number Genome 297) and Kyoto University (approval number G204).

#### Cell lines

HTLV-1-negative human T-cell lines (Jurkat [male] and Hut78 [male]) and ATL cell lines (TL-Om1 [male], ED [male], ATL55T+ [unspecified], MT1 [male] and HPB-ATL-2 [female]) were used in this study. They were cultured in RPMI-1640 supplemented with 10% fetal bovine serum (FBS). ATL55T+ was incubated with recombinant human interleukin 2 (rIL-2, 100 U/ml; PeproTech) because of its dependency. For HEK293 and HEK293T cells, DMEM was used with 10% FBS. The Platinum-E cell line, a retrovirus packaging cell line, was cultured in DMEM with 10% FBS, blasticidin (10 μg/mL; InvivoGen) and puromycin (1 μg/mL; InvivoGen). Stably HBZ-transduced lines (HBZ wild type [WT; the spliced form] or HBZ mutants) were generated from Jurkat cells with the Neon transfection system (Kinosada *et al*., 2017; Tanaka-Nakanishi et al., 2014). The HBZ WT and HBZ mutants were encoded into the pME18Sneo vector, which was modified from pCEV4.33 (Itoh et al., 1990). Three days after the electroporation, 1mg/mL G418 (Nacalai Tesque) was added and stable transfectants underwent selection for two weeks. All cell lines in this study were incubated at 37 C in a 5% CO_2_ humidified incubator.

#### Mice

C57BL/6J (CLEA Japan) and NOD/SCIDgc−/− (NSG; the Jackson laboratory) mice were purchased for the murine experiments in this study. The HBZ-transgenic mice (HBZ-Tg) contained the HBZ cDNA expressed under the murine CD4-specific promoter/enhancer/silencer, and were developed from the C57BL/6J mice as previously described (Satou *et al*., 2006). All HBZ-Tg mice were heterozygotes for the transgene. PCR-based genotyping for the HBZ-Tg mice was performed using following primers: HBZ-genotype-F: 5’-TGTAGGCTCAGATTCCCAACCAA-3’ and HBZ-genotype-R: 5’-TCTTCTCCTCAGCCCGTCGC-3’ as listed in Table S3. At sacrifice, primary splenocytes were collected from C57BL/6J mice and CD4^+^ T cells were then isolated with the Mouse CD4 T Lymphocyte Enrichment Set-DM (BD Biosciences).

#### Generation of TAp73−/− and DNp73−/− mice

The CRISPR/Cas9 system was utilized for isoform-specific knockout (KO) of the *Trp73* gene in C57BL/6J mice. Two types of small RNAs, target-recognizing CRISPR RNA (crRNA) and auxiliary trans-activating crRNA (tracrRNA), were purchased from FASMAC: TAp73-crRNA (5’-CAGAGAACUCCACAGGUGCUCGAAGG-3’), DNp73-crRNA (5’-CACGAGCCUACCAUGCUUUACGUCGG-3’) and tracrRNA (5’-AAACAGCAUAGCAAGUUAAAAUAAGGCUAGUCCGUUAUCAACUUGAAAAAGUGGCACCGAGUCGGUGCU-3’). Both crRNAs were designed with the CHOPCHOP web tool (Labun et al., 2019). Using electroporation, we delivered recombinant *Streptococcus pyogenes* Cas9 protein (TrueCut™ Cas9 Protein v2; Thermo Fisher Scientific) together with the crRNA and tracrRNA into in vitro fertilized eggs of C57BL/6J mice according to the institutional procedure of Kyoto university (Ishibashi et al., 2020). For electroporation, the Genome Editor (BEX) was used under conditions of 23 V (3 msec ON and 97 msec OFF) two times (Hashimoto et al., 2016). Only morphologically normal zygotes were microinjected at the pronuclear stage. Both TAp73 and DNp73 KO genotypes were obtained with 1bp insertions on the crRNA recognizing region, which results in truncating (frameshift) mutations (Figure S4A). Genotyping for each KO isoform was performed with restriction enzyme digestion of PCR-amplified DNA. For TAp73, genomic DNA was amplified with TAp73-genotype-F: 5’-GCCCTTCAGGACTATTAGAAAATGC-3’ and TAp73-genotype-R: 5’-GTAACCTGCATAACTGTGCATTACA-3’ (Table S3), and then BstBI (New England Biolabs) was used for cutting the PCR product. For DNp73, the genotype was similarly determined using PCR products amplified with DNp73-genotype-F: 5’-TGCTTGAGAGGAGCAACAGGGAG-3’ and DNp73-genotype-R: 5’-GGTGAGAATGCCAACTCTCAGT-3’ (Table S3), followed by MseI (New England Biolabs) enzymic restriction.

#### Murine xenograft model

2 × 10^6^ ED cells were subcutaneously inoculated in the right lower back of 6-week-old female NSG mice. Tumor growth was followed until the tumor reached a size of 40 mm^3^, at which point (Day 1), the NSG mice were randomly assigned to either the syrosingopine (7.5mg/kg; MedChemExpress) or the dimethyl sulfoxide (DMSO) group. Treatments were injected intraperitoneally every other day for a total of six doses. Tumor sizes were also measured with calipers every other day until Day 13. The xenograft model study was conducted under the condition that mice be euthanized if the tumor size exceeded 2000 mm^3^. At the end of the observation, the tumors were also weighed.

### METHOD DETAILS

#### Retroviral transduction of murine CD4^+^ T cells

Platinum-E cells were transduced with retroviral vectors containing the GFP and HBZ WT or HBZ mutant genes using Lipofectamine LTX with Plus Reagent (Thermo Fisher Scientific). The retroviruses were collected with the supernatant after 48 hours. To infect spleen-derived CD4^+^ T cells with the retrovirus, the remaining splenocytes were inactivated with 20 Gy of irradiation (150 kV and 20 mA) and used as antigen-presenting cells. CD4^+^ T cells were cultured with these antigen-presenting cells in the presence of anti-CD3 antibody (0.5 μg/mL; Mouse CD3 epsilon Antibody, R & D Systems), rIL-2 (100 U/ml; PeproTech) and 2-mercaptoethanol (55 μM) for 48 hours. The retroviral solution and polybrene (final concentration, 4mg/mL; Sigma-Aldrich) were then added and spin infection was performed at 3,000 rpm for 1 hour at room temperature. The transduced CD4^+^ T cells were then incubated at 37 C in 5% CO_2_ humidified incubator for 48 hours (Mitobe *et al*., 2015; Satou *et al*., 2011). After that, GFP-positive cells were collected for subsequent deep-sequencing.

#### RNA sequencing (RNA-seq)

Total RNA was collected from primary cells or cell lines with the ReliaPrep RNA Miniprep System (Promega). Libraries for RNA sequencing (RNA-seq) were generated with the TruSeq stranded mRNA Library Prep kit (Illumina). High-throughput sequencing was performed on the Illumina NovaSeq 6000 platform with a standard 100-bp paired-end read protocol.

#### Assay for transposase-accessible chromatin using sequencing (ATAC-seq)

Cells were incubated with Nextera Tn5 Transposase and 2x TD reaction buffer of the Illumina Tagment DNA Enzyme and Buffer Small Kit (Illumina) for 30 minutes at 37°C, according to the established procedure for ATAC-seq (Buenrostro et al., 2015). After purification with the Qiagen MinElute PCR Purification Kit, the transposed DNA was then amplified with the Nextera DNA Sample Preparation Kit (Illumina). Subsequently, the libraries were purified with the AMPure XP Reagent (Beckman Coulter) and quantified using the KAPA Library Quantification Kit (KAPA Biosystems). High-throughput sequencing was performed on the Illumina HiSeq 4000 platform with a standard 100-bp paired-end read protocol.

#### Reverse transcriptase quantitative-PCR (RT-qPCR)

Total RNA was purified using the ReliaPrep RNA Miniprep System (Promega) or TRIzol Reagent (Thermo Fisher Scientific) and then converted to cDNA with SuperScript IV (Thermo Fisher Scientific) according to the manufacturer’s instructions. Gene expression was assessed using FastStart Universal SYBR Green Master (Roche) on the StepOnePlus Real-Time PCR System (Applied Biosystems). Primers for RT-qPCR are listed in Table S3.

#### Immunoprecipitation and immunoblotting

Cell lysates were prepared with lysis buffer containing protease inhibitor cocktail (Nacalai Tesque) by incubating them for 30 minutes at 4°C, then centrifuged at 12,000rpm for 20 minutes. For immunoprecipitation, the lysates were incubated with primary antibody conjugated SureBeads Protein G Magnetic Beads (Bio-Rad Laboratories) for 1 hour rotating at room temperature. The immunocomplexes were then pulled out and washed 4 times with phosphate buffered saline with Tween-20 (Santa Cruz Biotechnology). For immunoblotting, the isolated precipitates or cell lysates were subjected to SDS-PAGE electrophoresis. For immunoprecipitation, antibodies against Flag (Sigma-Aldrich, #F3165), HA (MBL International, #M180-3), and HBZ, along with normal rabbit IgG, were used. The anti-HBZ antibody was produced in rabbit by immunization with HBZ peptides (CRGPPGEKAPPRGETH and QERRERKWRQGAEKC) as previously described (Higuchi et al., 2020). Immunoblotting was carried out with antibodies against HBZ described above, Flag (Sigma-Aldrich, #F7425), HA (MBL International, #561-5), α-Tubulin (Sigma-Aldrich, #T6199), EZH2 (Cell Signaling Technology, #5246S), TAp73 (Abcam, #ab14430) and DNp73 (Novus Biologicals, #NBP2-24873). These antibodies were then detected with the appropriate secondary antibody conjugated to horseradish peroxidase (HRP; Thermo Fisher Scientific, #31430 for mouse and #G-21234 for rabbit). Immunoblots were visualized with the ChemiDoc imaging system (Bio-Rad Laboratories).

#### Chromatin immunoprecipitation (ChIP)

ChIP assays were conducted with the use of SimpleChIP Enzymatic Chromatin IP Kit (Magnetic Beads; Cell Signaling Technology) according to the manufacturer’s instructions. Briefly, cross-linked chromatin prepared from 4 × 10^6^ cells was immunoprecipitated overnight with antibodies against EZH2 (Cell Signaling Technology, #5246S), H3K27me3 (Cell Signaling Technology, #9733S), IRF4 (Cell Signaling Technology, #4964S), BATF3 (R & D Systems, #AF7437) or TAp73 (Abcam, # ab14430). The immunocomplexes were then conjugated with ChIP-Grade Protein G Magnetic Beads (Cell Signaling Technology) for 2 hours at 4°C. After 4 washes, the bead-bound DNA was eluted by gentle vortexing (1,200 rpm) for 30 minutes at 65°C. Purified DNA was quantified by ChIP-qPCR in comparison to a 2% input control. The primers for ChIP-qPCR are listed in Table S3. The eluted DNAs and the input were also subjected to ChIP sequencing (ChIP-seq). The TruSeq ChIP Library Preparation Kit (Illumina) was used for library preparation. Libraries were sequenced on the Illumina NovaSeq6000 platform with a standard 150-bp paired-end read protocol.

#### Histone methyltransferase (HMT) activity assay

Using the EpiQuik Nuclear Extraction Kit (EpigenTek), we isolated nuclear proteins from Jurkat cells that were stably transduced with WT or mutant HBZ. The nuclear extracts were then subjected to assays of histone methyltransferase activity that specifically target histone H3 at lysine 27. We used the EpiQuik Histone Methyltransferase Activity/Inhibition Assay Kit (H3-K27) (EpigenTek). We measured the absorbance at 450 nm for each sample and generated a standard curve. The HMT activity (OD/h/mg) was calculated according to the manufacturer’s instructions.

#### Luciferase assay

The vectors for the luciferase assay, pGL4.10[luc2] (#E6651) and pNL3.2.CMV (#N141A), were purchased from Promega. The latter was edited for each promoter assay. For Figure 3D, the hg19 genome region of chr1:3593076-3594185 was cloned using the Zero Blunt Cloning Kit (Thermo Fisher Scientific) and inserted between the Kpn1 and Xho1 sites of pNL3.2.CMV. For all remaining luciferase assays in this study, the minimal promoter (minP) was removed using the In-Fusion Snap Assembly Master Mix (TaKaRa), and the appropriate sequences (enumerated in Table S3) were inserted between the Kpn1 and Xho1 sites of pNL3.2.CMV. Ectopic gene expression in the luciferase study was induced by pME18S or pCAGGS containing the corresponding coding sequence. HEK293 or Jurkat cells were seeded into a 24-well plate at 1 × 10^5^ cells per well. After 24 hours, cells were transfected with the indicated plasmids using TransIT (for HEK293) or the Neon transfection system (for Jurkat). After 24 hours incubation, the transfected cells were lysed in Passive Lysis 5X Buffer (Promega). Firefly luciferase (FLuc) and NanoLuc luciferase (NLuc) were detected by the Nano-Glo Dual-Luciferase Reporter Assay System (Promega) and measured using a TriStar LB941 and MikroWin 2000 version 4.41 (Berthold Technologies). Relative luminescence was calculated as NLuc per FLuc.

#### Lentiviral transduction of ATL cell lines

For generation of lentiviral particles, we transduced psPAX2, pCMV-VSV-G and shRNA-encoding lentiviral vectors (Sigma-Aldrich) into HEK293T cells with Lipofectamine 2000 Transfection Reagent (Thermo Fisher Scientific). For subsequent GFP competition assays, the MISSION TRC2 pLKO.5-puro plasmid (Sigma-Aldrich) was modified to replace the puromycin-resistance gene with the enhanced green fluorescent protein (EGFP) gene using the In-Fusion Snap Assembly Master Mix (TaKaRa). Forty-eight hours after transfection, the supernatant was collected and ultracentrifuged (25,000 rpm) for 2 hours at 4°C. Enriched lentiviruses were infected into ATL cell lines using RetroNectin (TaKaRa) with spin infection (3,000rpm) for 1 hour.

#### Flow cytometry and GFP competition assay

After lentiviral transduction (counted as Day 1), GFP positivity of ATL cell lines was analyzed by flow cytometer (FACSVerse) every other day from Day 4 to 12. GFP was evaluated only in live cells. The GFP-positive rate on Day 4 was used as a baseline to evaluate changes in positivity rate. The cells were analyzed using the LIVE/DEAD Fixable Dead Cell Stain Kit (Thermo Fisher Scientific), Alexa Fluor 647 Annexin V (BioLegend) and pHrodo Red AM Intracellular pH Indicator (Thermo Fisher Scientific). Extracted data were analyzed by the FlowJo software (BD Bioscience).

#### Lactate assay

The Lactate Assay Kit-WST (Dojindo) was used according to the manufacturer’s instructions. Intracellular lactate was assayed after lysis of cells with 0.1% Triton X-100 (Nacalai Tesque). After incubation at 37°C for 30 minutes, the absorbance at 450 nm was measured by using the TriStar LB941 and MikroWin 2000 version 4.41 (Berthold Technologies). In experiments in which ATL cell lines were treated with syrosingopine, extracellular and intracellular lactate were measured after treatment with syrosingopine for 4 days, respectively.

#### Metabolomic analysis by gas chromatography-mass spectrometry (GC-MS)

Isolated murine CD4^+^ T-cell pellets were suspended in 250 μl of 100% methanol and frozen in liquid nitrogen. After sonication, the samples were centrifuged at 15,000 rpm at 4°C for 15 minutes. Subsequently, the upper aqueous layer was transferred to a new tube, mixed with 250 μL of Milli-Q water and shaken at 1,500 rpm at 4°C for 15 minutes. The solution was dried using a Thermo Scientific Savant DNA SpeedVac Concentrator Kit (DNA120OP230, Thermo Fisher Scientific) and then lyophilized with the FTS Systems Flexi-Dry Freeze Dryer Model (FD-1-84A, FTS). After derivatization with O-Methylhydroxylamine Hydrochloride/pyridine (Tokyo Chemical Industry) and N-methyl-N-trimethylsilyltrifluoroacetamide (GL Sciences), the samples were analyzed on a Shimadzu TQ8050 triple quadrupole mass spectrometer (Shimadzu). The chromatograms and mass spectra were analyzed with GC-MS solution software (Shimadzu). Compounds were identified using the Smart Metabolites Database (Shimadzu). The data obtained were then standardized and analyzed using MetaboAnalyst (Chong et al., 2018).

#### Metabolic flux assay

After coating Seahorse XFe24 Cell Culture Microplates (Agilent) with Poly-D-lysine hydrobromide (FUJIFILM Wako Pure Chemical), we seeded isolated mouse CD4^+^ T cells at 1 × 10^6^ cells per well. The cells were incubated in Seahorse XF RPMI medium, pH 7.4 (Agilent) supplemented with 2 mM Seahorse XF Glutamine Solution (Agilent) at 37°C. We used the Seahorse XF Glycolysis Stress Test Kit (Agilent) according to the manufacturer’s instructions. The assay was carried out on the Seahorse XFe24 Analyzer (Agilent). Extracted data were analyzed using Seahorse Wave Desktop Software (Agilent).

#### Cell proliferation and viability assay

Cell lines were treated with syrosingopine (MedChemExpress, an MCT1/4 inhibitor). At first, syrosingopine was dissolved in DMSO. The dissolved syrosingopine or DMSO alone was finally diluted 100-fold in RPMI medium with cell lines, and used for further experiments. Cells were then seeded into a 96-well plate without removing syrosingopine and DMSO throughout the experiments. To assess cell growth, 20 μL of 3-(4,5-dimethylthiazol-2-yl)-2,5-diphenyltetrazolium bromide (MTT) was added to each well. Cells were kept in the dark in a humidified incubator at 37°C with 5% CO_2_. Two hours later, cells were lysed with 100μL of lysis buffer (4% Triton X-100, and 0.14% HCl in 2-propanol). Then the absorbance at 450nm was measured by using a TriStar LB941 and MikroWin 2000 version 4.41 (Berthold Technologies). The MTT assay was started 3 hours after the seeding (counted as day 0). Cell viability was assessed by the Countess Automated Cell Counter (Thermo Fisher Scientific) at day 4.

### QUANTIFICATION AND STATISTICAL ANALYSIS

#### Analysis of RNA-seq

In addition to the sequencing data in this study, RNA-seq data were obtained from the following public archives: GSE74246 (Corces et al., 2016) and GSE162712 (Bauer et al., 2021) for normal CD4^+^ T cells; and SRP042199 (Nakagawa et al., 2014), GSE143986 (Yoshida et al., 2020) and DRP006659 (Tanaka et al., 2020) for ATL cells. After fastq files were cleansed with Trim Galore, obtained reads were mapped to the hg19 or mm10 genome sequence. To align them, HISAT2 (Kim et al., 2019) was used with the following options: --dta-cufflinks, --no-discordant, --no-mixed and --rna- strandness FR. The aligned data were then sorted by samtools (Danecek et al., 2021). After counting reads in features with htseq-count of HTSeq (Putri et al., 2022), differentially expressed genes were identified if the fold change was greater than 1.1 and the adjusted p-value (padj) was less than 0.05 in the DESeq2 analysis (Love et al., 2014). The results were then subjected to Gene Set Enrichment Analysis (GSEA) (Subramanian et al., 2005) using clusterProfiler (Wu et al., 2021). Transcripts per million (TPM) were calculated using TPMCalculator (Vera Alvarez et al., 2019).

#### Cleansing and mapping of DNA sequencing data (ATAC-seq and ChIP-seq)

As with the RNA-seq analysis, fastq files of both ATAC-seq and ChIP-seq data were trimmed and qualified by the Trim Galore. Mapping to the corresponding genome was carried out by Bowtie2 with the “--very-sensitive” option (Langmead and Salzberg, 2012). To remove mitochondrial reads and reads which aligned twice or more, we removed “MT” and/or “XS” tagged reads using samtools. Subsequently, reads with a mapping quality of more than 30 were selected using samtools view with the “-q 30” option. BAM files were also cleaned up by removing PCR duplicates with Picard (MarkDuplicates) and the ENCODE Blacklist genome regions (Amemiya et al., 2019) using BEDtools (intersectBed) (Quinlan and Hall, 2010).

#### Peak calling of ATAC-seq data in retrovirally HBZ-transduced mice samples

MACS2 (Zhang et al., 2008) was used for peak calling with the following options: “-- nomodel --nolambda --keep-dup all --call-summits”. We further used HOMER (Heinz et al., 2010) for subsequent analysis including motif analysis. Peak bed files from the MACS2 results were converted to a HOMER bed file using the bed2pos.pl tool. After merging the peaks with mergePeaks, we counted each extracted peak using HTSeq. Using these counts with DESeq2, we compared samples from cells transduced with WT HBZ. mutant HBZ or vector alone. Statistically significant peaks (defined as peaks with padj less than 0.05) were subjected to motif analysis using findMotifsGenome.pl of HOMER with the “-size 200 -mask” options.

#### Peak calling of ChIP-seq data

Using the corresponding input sequenced file as a control, ChIP-seq peaks were assessed by MACS2 (Zhang *et al*., 2008) with the “-p 1e-3” option. Extracted peaks were then annotated to the nearest genes of the hg19 by using the Genomic Regions Enrichment of Annotations Tool (GREAT) (McLean et al., 2010). We also utilized publicly available ChIP-seq databases via the ChIP-Atlas (Zou et al., 2022) with the following ID: CD4^+^ T cells of HBZ-Tg mice for H3K27ac (DRP002840; DRX028570) (Yasuma *et al*., 2016), TL-Om1 cells for H3K27ac (GSE85692; SRX2026362) (Wong et al., 2017), murine CD4^+^ Th2 cells for IRF4 and BATF3 (GEO: GSE85172; SRX2646182 and SRX2646184) (Iwata et al., 2017) and KK1 cells for IRF4 and BATF3 (GEO: GSE94732; SRX2548278 and SRX2548284) (Nakagawa *et al*., 2018). For generating the heatmaps of H3K27me3 genomic distribution, we used deepTools (Ramírez et al., 2016). First, we extracted peaks using the bamCompare tool with the “-- operation subtract” option. To calculate enriched scores per genome regions for each transcript, the bigWig files were then analyzed using the computeMatrix tool (reference- point mode) with the “--beforeRegionStartLength 3000 --regionBodyLength 5000 -- afterRegionStartLength 3000 --skipZeros” options. Subsequently, heatmaps were generated using the plotHeatmap tool with the “--kmeans 5 --sortRegions=descend” options. The cluster 2 genes (Tables S1 and S2) were subjected to KEGG pathway analysis by using the clusterProfiler. For RNA-seq and ChIP-seq combined scatter plots, we first calculated the AUC of cluster 2 (4,297 genes) from ChIP-seq (H3K27me3 and EZH2) data of the WT sample by using the computeMatrix tool with the “-- referencePoint TSS -a 5000 -b 5000. Then, the AUC values were combined with fold changes calculated from RNA-seq data.

#### Analysis of TCGA data

Raw data were obtained from the official website of TCGA Pan-Cancer (PANCAN). We analyzed the following datasets: (1) tcga_RSEM_gene_tpm, (2) tcga_rsem_isoform_tpm and (3) TCGA_survival_data_2.txt. We also utilized a web server called GEPIA2 to analyze and visualize TCGA data (Tang et al., 2019). The abbreviations used in TCGA are found in the following website: (https://gdc.cancer.gov/resources-tcga-users/tcga-code-tables/tcga-study-abbreviations).

#### Statistical Analysis and data visualization

The student’s t-test (two-sided) was used for comparisons between the two groups, and differences were defined as statistically significant for p < 0.05. For intercomparison of more than 2 groups, a one-way ANOVA followed by a post-hoc test (Turkey, Dunnet [parametric] or Steel [non-parametric]) was applied. To assess the association between targeted gene expressions, we used Pearson correlation analysis. In mouse experiments, the sample size was not pre-selected and no inclusion/exclusion criteria were used. The cumulative incidence of inflammation was analyzed by Gray test. The probability of survival was estimated by the Kaplan-Meier method, and hazard ratios (HR) were estimated with the Cox regression model. All analyses were conducted using R Statistical Software (v4.1.2). The bar and box plots contain the mean value and standard deviation calculated from at least three biological replicates. To generate these plots, we used ggplot2 and pheatmap. To visualize deep-sequencing data, we used deepTools (Ramírez *et al*., 2016) and SparK (Kurtenbach and William Harbour, 2019).

## Reference

Akkouche, A., Moodad, S., Hleihel, R., Skayneh, H., Chambeyron, S., El Hajj, H., and Bazarbachi, A. (2021). In vivo antagonistic role of the Human T-Cell Leukemia Virus Type 1 regulatory proteins Tax and HBZ. PLoS Pathog 17, e1009219. 10.1371/journal.ppat.1009219.

Amemiya, H.M., Kundaje, A., and Boyle, A.P. (2019). The ENCODE Blacklist: Identification of Problematic Regions of the Genome. Scientific Reports 9, 9354. 10.1038/s41598-019-45839-z.

Bauer, L., Müller, L.J., Volkers, S.M., Heinrich, F., Mashreghi, M.F., Ruppert, C., Sander, L.E., and Hutloff, A. (2021). Follicular Helper-like T Cells in the Lung Highlight a Novel Role of B Cells in Sarcoidosis. Am J Respir Crit Care Med 204, 1403–1417. 10.1164/rccm.202012-4423OC.

Benjamin, D., Colombi, M., Hindupur, S.K., Betz, C., Lane, H.A., El-Shemerly, M.Y., Lu, M., Quagliata, L., Terracciano, L., Moes, S., et al. (2016). Syrosingopine sensitizes cancer cells to killing by metformin. Sci Adv 2, e1601756. 10.1126/sciadv.1601756.

Benjamin, D., Robay, D., Hindupur, S.K., Pohlmann, J., Colombi, M., El-Shemerly, M.Y., Maira, S.M., Moroni, C., Lane, H.A., and Hall, M.N. (2018). Dual Inhibition of the Lactate Transporters MCT1 and MCT4 Is Synthetic Lethal with Metformin due to NAD+ Depletion in Cancer Cells. Cell Rep 25, 3047–3058.e3044. 10.1016/j.celrep.2018.11.043.

Buenrostro, J.D., Wu, B., Chang, H.Y., and Greenleaf, W.J. (2015). ATAC-seq: A Method for Assaying Chromatin Accessibility Genome-Wide. Curr Protoc Mol Biol 109, 21.29.21–21.29.29. 10.1002/0471142727.mb2129s109.

Buxbaum, A.R., Haimovich, G., and Singer, R.H. (2015). In the right place at the right time: visualizing and understanding mRNA localization. Nat Rev Mol Cell Biol 16, 95–109. 10.1038/nrm3918.

Candeias, M.M., Malbert-Colas, L., Powell, D.J., Daskalogianni, C., Maslon, M.M., Naski, N., Bourougaa, K., Calvo, F., and Fåhraeus, R. (2008). P53 mRNA controls p53 activity by managing Mdm2 functions. Nat Cell Biol 10, 1098–1105. 10.1038/ncb1770.

Chong, J., Soufan, O., Li, C., Caraus, I., Li, S., Bourque, G., Wishart, D.S., and Xia, J. (2018). MetaboAnalyst 4.0: towards more transparent and integrative metabolomics analysis. Nucleic Acids Res 46, W486–w494. 10.1093/nar/gky310.

Corces, M.R., Buenrostro, J.D., Wu, B., Greenside, P.G., Chan, S.M., Koenig, J.L., Snyder, M.P., Pritchard, J.K., Kundaje, A., Greenleaf, W.J., et al. (2016). Lineage-specific and single-cell chromatin accessibility charts human hematopoiesis and leukemia evolution. Nat Genet 48, 1193–1203. 10.1038/ng.3646.

Danecek, P., Bonfield, J.K., Liddle, J., Marshall, J., Ohan, V., Pollard, M.O., Whitwham, A., Keane, T., McCarthy, S.A., Davies, R.M., and Li, H. (2021). Twelve years of SAMtools and BCFtools. GigaScience 10. 10.1093/gigascience/giab008.

Davis, C.A., Hitz, B.C., Sloan, C.A., Chan, E.T., Davidson, J.M., Gabdank, I., Hilton, J.A., Jain, K., Baymuradov, U.K., Narayanan, A.K., et al. (2018). The Encyclopedia of DNA elements (ENCODE): data portal update. Nucleic Acids Res 46, D794–d801. 10.1093/nar/gkx1081.

DeBerardinis, R.J., Lum, J.J., Hatzivassiliou, G., and Thompson, C.B. (2008). The biology of cancer: metabolic reprogramming fuels cell growth and proliferation. Cell Metab 7, 11–20. 10.1016/j.cmet.2007.10.002.

Du, W., Jiang, P., Mancuso, A., Stonestrom, A., Brewer, M.D., Minn, A.J., Mak, T.W., Wu, M., and Yang, X. (2013). TAp73 enhances the pentose phosphate pathway and supports cell proliferation. Nat Cell Biol 15, 991–1000. 10.1038/ncb2789.

ENCODE.Project.Consortium (2012). An integrated encyclopedia of DNA elements in the human genome. Nature 489, 57–74. 10.1038/nature11247.

Engelmann, D., Meier, C., Alla, V., and Pützer, B.M. (2015). A balancing act: orchestrating amino-truncated and full-length p73 variants as decisive factors in cancer progression. Oncogene 34, 4287–4299. 10.1038/onc.2014.365.

Fujikawa, D., Nakagawa, S., Hori, M., Kurokawa, N., Soejima, A., Nakano, K., Yamochi, T., Nakashima, M., Kobayashi, S., Tanaka, Y., et al. (2016). Polycomb-dependent epigenetic landscape in adult T-cell leukemia. Blood 127, 1790–1802. 10.1182/blood-2015-08-662593.

Gazon, H., Chauhan, P.S., Porquet, F., Hoffmann, G.B., Accolla, R., and Willems, L. (2020). Epigenetic silencing of HTLV-1 expression by the HBZ RNA through interference with the basal transcription machinery. Blood advances 4, 5574–5579. 10.1182/bloodadvances.2020001675.

Grob, T.J., Novak, U., Maisse, C., Barcaroli, D., Lüthi, A.U., Pirnia, F., Hügli, B., Graber, H.U., De Laurenzi, V., Fey, M.F., et al. (2001). Human ΔNp73 regulates a dominant negative feedback loop for TAp73 and p53. Cell Death & Differentiation 8, 1213–1223. 10.1038/sj.cdd.4400962.

Hanahan, D. (2022). Hallmarks of Cancer: New Dimensions. Cancer discovery 12, 31–46. 10.1158/2159-8290.Cd-21-1059.

Hanahan, D., and Weinberg, R.A. (2011). Hallmarks of cancer: the next generation. Cell 144, 646–674. 10.1016/j.cell.2011.02.013.

Hashimoto, M., Yamashita, Y., and Takemoto, T. (2016). Electroporation of Cas9 protein/sgRNA into early pronuclear zygotes generates non-mosaic mutants in the mouse. Dev Biol 418, 1–9. 10.1016/j.ydbio.2016.07.017.

Heinz, S., Benner, C., Spann, N., Bertolino, E., Lin, Y.C., Laslo, P., Cheng, J.X., Murre, C., Singh, H., and Glass, C.K. (2010). Simple combinations of lineage-determining transcription factors prime cis-regulatory elements required for macrophage and B cell identities. Mol Cell 38, 576–589. 10.1016/j.molcel.2010.05.004.

Higuchi, Y., Yasunaga, J.I., Mitagami, Y., Tsukamoto, H., Nakashima, K., Ohshima, K., and Matsuoka, M. (2020). HTLV-1 induces T cell malignancy and inflammation by viral antisense factor-mediated modulation of the cytokine signaling. Proceedings of the National Academy of Sciences of the United States of America 117, 13740–13749. 10.1073/pnas.1922884117.

Hubé, F., Velasco, G., Rollin, J., Furling, D., and Francastel, C. (2010). Steroid receptor RNA activator protein binds to and counteracts SRA RNA-mediated activation of MyoD and muscle differentiation. Nucleic Acids Research 39, 513–525. 10.1093/nar/gkq833.

Ishibashi, R., Abe, K., Ido, N., Kitano, S., Miyachi, H., and Toyoshima, F. (2020). Genome editing with the donor plasmid equipped with synthetic crRNA-target sequence. Sci Rep 10, 14120. 10.1038/s41598-020-70804-6.

Ishio, T., Kumar, S., Shimono, J., Daenthanasanmak, A., Dubois, S., Lin, Y., Bryant, B., Petrus, M.N., Bachy, E., Huang, D.W., et al. (2022). Genome-wide CRISPR screen identifies CDK6 as a therapeutic target in adult T-cell leukemia/lymphoma. Blood 139, 1541–1556. 10.1182/blood.2021012734.

Itoh, N., Yonehara, S., Schreurs, J., Gorman, D.M., Maruyama, K., Ishii, A., Yahara, I., Arai, K., and Miyajima, A. (1990). Cloning of an interleukin-3 receptor gene: a member of a distinct receptor gene family. Science 247, 324–327. 10.1126/science.2404337.

Iwata, A., Durai, V., Tussiwand, R., Briseño, C.G., Wu, X., Grajales-Reyes, G.E., Egawa, T., Murphy, T.L., and Murphy, K.M. (2017). Quality of TCR signaling determined by differential affinities of enhancers for the composite BATF-IRF4 transcription factor complex. Nature immunology 18, 563–572. 10.1038/ni.3714.

Jones, R.G., and Thompson, C.B. (2009). Tumor suppressors and cell metabolism: a recipe for cancer growth. Genes Dev 23, 537–548. 10.1101/gad.1756509.

Kim, D., Paggi, J.M., Park, C., Bennett, C., and Salzberg, S.L. (2019). Graph-based genome alignment and genotyping with HISAT2 and HISAT-genotype. Nature Biotechnology 37, 907–915. 10.1038/s41587-019-0201-4.

Kinosada, H., Yasunaga, J.I., Shimura, K., Miyazato, P., Onishi, C., Iyoda, T., Inaba, K., and Matsuoka, M. (2017). HTLV-1 bZIP Factor Enhances T-Cell Proliferation by Impeding the Suppressive Signaling of Co-inhibitory Receptors. PLoS Pathog 13, e1006120. 10.1371/journal.ppat.1006120.

Kogure, Y., Kameda, T., Koya, J., Yoshimitsu, M., Nosaka, K., Yasunaga, J.I., Imaizumi, Y., Watanabe, M., Saito, Y., Ito, Y., et al. (2021). Whole-genome landscape of adult T-cell leukemia/lymphoma. Blood. 10.1182/blood.2021013568.

Koppenol, W.H., Bounds, P.L., and Dang, C.V. (2011). Otto Warburg’s contributions to current concepts of cancer metabolism. Nat Rev Cancer 11, 325–337. 10.1038/nrc3038.

Kurtenbach, S., and William Harbour, J. (2019). SparK: A Publication-quality NGS Visualization Tool. bioRxiv, 845529. 10.1101/845529.

Labun, K., Montague, T.G., Krause, M., Torres Cleuren, Y.N., Tjeldnes, H., and Valen, E. (2019). CHOPCHOP v3: expanding the CRISPR web toolbox beyond genome editing. Nucleic Acids Research 47, W171–W174. 10.1093/nar/gkz365.

Langmead, B., and Salzberg, S.L. (2012). Fast gapped-read alignment with Bowtie 2. Nat Methods 9, 357–359. 10.1038/nmeth.1923.

Laugesen, A., and Helin, K. (2014). Chromatin repressive complexes in stem cells, development, and cancer. Cell stem cell 14, 735–751. 10.1016/j.stem.2014.05.006.

Li, L., Li, L., Li, W., Chen, T., Bin, Z., Zhao, L., Wang, H., Wang, X., Xu, L., Liu, X., et al. (2018). TAp73-induced phosphofructokinase-1 transcription promotes the Warburg effect and enhances cell proliferation. Nature communications 9, 4683. 10.1038/s41467-018-07127-8.

Love, M.I., Huber, W., and Anders, S. (2014). Moderated estimation of fold change and dispersion for RNA-seq data with DESeq2. Genome Biology 15, 550. 10.1186/s13059-014-0550-8.

Ma, G., Yasunaga, J., and Matsuoka, M. (2016). Multifaceted functions and roles of HBZ in HTLV-1 pathogenesis. Retrovirology 13, 16. 10.1186/s12977-016-0249-x.

Ma, G., Yasunaga, J.I., Shimura, K., Takemoto, K., Watanabe, M., Amano, M., Nakata, H., Liu, B., Zuo, X., and Matsuoka, M. (2021). Human retroviral antisense mRNAs are retained in the nuclei of infected cells for viral persistence. Proceedings of the National Academy of Sciences of the United States of America 118. 10.1073/pnas.2014783118.

Matsuoka, M., and Jeang, K.T. (2007). Human T-cell leukaemia virus type 1 (HTLV-1) infectivity and cellular transformation. Nat Rev Cancer 7, 270–280. 10.1038/nrc2111.

Matsuoka, M., and Mesnard, J.M. (2020). HTLV-1 bZIP factor: the key viral gene for pathogenesis. Retrovirology 17, 2. 10.1186/s12977-020-0511-0.

McLean, C.Y., Bristor, D., Hiller, M., Clarke, S.L., Schaar, B.T., Lowe, C.B., Wenger, A.M., and Bejerano, G. (2010). GREAT improves functional interpretation of cis-regulatory regions. Nat Biotechnol 28, 495–501. 10.1038/nbt.1630.

Melino, G., De Laurenzi, V., and Vousden, K.H. (2002). p73: Friend or foe in tumorigenesis. Nature Reviews Cancer 2, 605–615. 10.1038/nrc861.

Mesri, E.A., Feitelson, M.A., and Munger, K. (2014). Human viral oncogenesis: a cancer hallmarks analysis. Cell Host Microbe 15, 266–282. 10.1016/j.chom.2014.02.011.

Mitobe, Y., Yasunaga, J., Furuta, R., and Matsuoka, M. (2015). HTLV-1 bZIP Factor RNA and Protein Impart Distinct Functions on T-cell Proliferation and Survival. Cancer Res 75, 4143–4152. 10.1158/0008-5472.Can-15-0942.

Nakagawa, M., Schmitz, R., Xiao, W., Goldman, C.K., Xu, W., Yang, Y., Yu, X., Waldmann, T.A., and Staudt, L.M. (2014). Gain-of-function CCR4 mutations in adult T cell leukemia/lymphoma. J Exp Med 211, 2497–2505. 10.1084/jem.20140987.

Nakagawa, M., Shaffer, A.L., 3rd, Ceribelli, M., Zhang, M., Wright, G., Huang, D.W., Xiao, W., Powell, J., Petrus, M.N., Yang, Y., et al. (2018). Targeting the HTLV-I- Regulated BATF3/IRF4 Transcriptional Network in Adult T Cell Leukemia/Lymphoma. Cancer Cell 34, 286–297.e210. 10.1016/j.ccell.2018.06.014.

Nakahata, S., Ichikawa, T., Maneesaay, P., Saito, Y., Nagai, K., Tamura, T., Manachai, N., Yamakawa, N., Hamasaki, M., Kitabayashi, I., et al. (2014). Loss of NDRG2 expression activates PI3K-AKT signalling via PTEN phosphorylation in ATLL and other cancers. Nature communications 5, 3393. 10.1038/ncomms4393.

Ong, J.Z.L., Yokomori, R., Wong, R.W.J., Tan, T.K., Ueda, R., Ishida, T., Iida, S., and Sanda, T. (2022). Requirement for TP73 and genetic alterations originating from its intragenic super-enhancer in adult T-cell leukemia. Leukemia. 10.1038/s41375-022-01655-5.

Putri, G.H., Anders, S., Pyl, P.T., Pimanda, J.E., and Zanini, F. (2022). Analysing high-throughput sequencing data in Python with HTSeq 2.0. Bioinformatics (Oxford, England) 38, 2943–2945. 10.1093/bioinformatics/btac166.

Quinlan, A.R., and Hall, I.M. (2010). BEDTools: a flexible suite of utilities for comparing genomic features. Bioinformatics (Oxford, England) 26, 841–842. 10.1093/bioinformatics/btq033.

Rabinowitz, J.D., and Enerbäck, S. (2020). Lactate: the ugly duckling of energy metabolism. Nat Metab 2, 566–571. 10.1038/s42255-020-0243-4.

Ramírez, F., Ryan, D.P., Grüning, B., Bhardwaj, V., Kilpert, F., Richter, A.S., Heyne, S., Dündar, F., and Manke, T. (2016). deepTools2: a next generation web server for deep-sequencing data analysis. Nucleic Acids Res 44, W160–165. 10.1093/nar/gkw257.

Sampath, K., and Ephrussi, A. (2016). CncRNAs: RNAs with both coding and non-coding roles in development. Development 143, 1234–1241. 10.1242/dev.133298.

Sasaki, D., Imaizumi, Y., Hasegawa, H., Osaka, A., Tsukasaki, K., Choi, Y.L., Mano, H., Marquez, V.E., Hayashi, T., Yanagihara, K., et al. (2011). Overexpression of Enhancer of zeste homolog 2 with trimethylation of lysine 27 on histone H3 in adult T-cell leukemia/lymphoma as a target for epigenetic therapy. Haematologica 96, 712–719. 10.3324/haematol.2010.028605.

Satou, Y., Yasunaga, J., Yoshida, M., and Matsuoka, M. (2006). HTLV-I basic leucine zipper factor gene mRNA supports proliferation of adult T cell leukemia cells. Proceedings of the National Academy of Sciences of the United States of America 103, 720–725. 10.1073/pnas.0507631103.

Satou, Y., Yasunaga, J., Zhao, T., Yoshida, M., Miyazato, P., Takai, K., Shimizu, K., Ohshima, K., Green, P.L., Ohkura, N., et al. (2011). HTLV-1 bZIP factor induces T-cell lymphoma and systemic inflammation in vivo. PLoS Pathog 7, e1001274. 10.1371/journal.ppat.1001274.

Stiewe, T., and Pützer, B.M. (2000). Role of the p53-homologue p73 in E2F1-induced apoptosis. Nat Genet 26, 464–469. 10.1038/82617.

Subramanian, A., Tamayo, P., Mootha, V.K., Mukherjee, S., Ebert, B.L., Gillette, M.A., Paulovich, A., Pomeroy, S.L., Golub, T.R., Lander, E.S., and Mesirov, J.P. (2005). Gene set enrichment analysis: A knowledge-based approach for interpreting genome-wide expression profiles. Proceedings of the National Academy of Sciences 102, 15545–15550. doi:10.1073/pnas.0506580102.

Sugata, K., Yasunaga, J., Kinosada, H., Mitobe, Y., Furuta, R., Mahgoub, M., Onishi, C., Nakashima, K., Ohshima, K., and Matsuoka, M. (2016). HTLV-1 Viral Factor HBZ Induces CCR4 to Promote T-cell Migration and Proliferation. Cancer Res 76, 5068–5079. 10.1158/0008-5472.Can-16-0361.

Tanaka, A., Ishitsuka, Y., Ohta, H., Fujimoto, A., Yasunaga, J.I., and Matsuoka, M. (2020). Systematic clustering algorithm for chromatin accessibility data and its application to hematopoietic cells. PLoS computational biology 16, e1008422. 10.1371/journal.pcbi.1008422.

Tanaka-Nakanishi, A., Yasunaga, J., Takai, K., and Matsuoka, M. (2014). HTLV-1 bZIP factor suppresses apoptosis by attenuating the function of FoxO3a and altering its localization. Cancer Res 74, 188–200. 10.1158/0008-5472.Can-13-0436.

Tang, Z., Kang, B., Li, C., Chen, T., and Zhang, Z. (2019). GEPIA2: an enhanced web server for large-scale expression profiling and interactive analysis. Nucleic Acids Research 47, W556–W560. 10.1093/nar/gkz430.

Tomasini, R., Tsuchihara, K., Wilhelm, M., Fujitani, M., Rufini, A., Cheung, C.C., Khan, F., Itie-Youten, A., Wakeham, A., Tsao, M.S., et al. (2008). TAp73 knockout shows genomic instability with infertility and tumor suppressor functions. Genes Dev 22, 2677–2691. 10.1101/gad.1695308.

Toyoda, K., and Matsuoka, M. (2022). Functional and Pathogenic Roles of Retroviral Antisense Transcripts. Front Immunol 13, 875211. 10.3389/fimmu.2022.875211.

Vera Alvarez, R., Pongor, L.S., Mariño-Ramírez, L., and Landsman, D. (2019). TPMCalculator: one-step software to quantify mRNA abundance of genomic features. Bioinformatics (Oxford, England) 35, 1960–1962. 10.1093/bioinformatics/bty896.

Warburg, O. (1956). On the origin of cancer cells. Science 123, 309–314. 10.1126/science.123.3191.309.

Wong, R.W.J., Ngoc, P.C.T., Leong, W.Z., Yam, A.W.Y., Zhang, T., Asamitsu, K., Iida, S., Okamoto, T., Ueda, R., Gray, N.S., et al. (2017). Enhancer profiling identifies critical cancer genes and characterizes cell identity in adult T-cell leukemia. Blood 130, 2326–2338. 10.1182/blood-2017-06-792184.

Wu, T., Hu, E., Xu, S., Chen, M., Guo, P., Dai, Z., Feng, T., Zhou, L., Tang, W., Zhan, L., et al. (2021). clusterProfiler 4.0: A universal enrichment tool for interpreting omics data. The Innovation 2. 10.1016/j.xinn.2021.100141.

Yamagishi, M., Fujikawa, D., Watanabe, T., and Uchimaru, K. (2018). HTLV-1-Mediated Epigenetic Pathway to Adult T-Cell Leukemia-Lymphoma. Front Microbiol 9, 1686. 10.3389/fmicb.2018.01686.

Yamagishi, M., Hori, M., Fujikawa, D., Ohsugi, T., Honma, D., Adachi, N., Katano, H., Hishima, T., Kobayashi, S., Nakano, K., et al. (2019). Targeting Excessive EZH1 and EZH2 Activities for Abnormal Histone Methylation and Transcription Network in Malignant Lymphomas. Cell Rep 29, 2321–2337.e2327. 10.1016/j.celrep.2019.10.083.

Yamagishi, M., Nakano, K., Miyake, A., Yamochi, T., Kagami, Y., Tsutsumi, A., Matsuda, Y., Sato-Otsubo, A., Muto, S., Utsunomiya, A., et al. (2012). Polycomb-mediated loss of miR-31 activates NIK-dependent NF-κB pathway in adult T cell leukemia and other cancers. Cancer Cell 21, 121–135. 10.1016/j.ccr.2011.12.015.

Yan, X., Hoek, T.A., Vale, R.D., and Tanenbaum, M.E. (2016). Dynamics of Translation of Single mRNA Molecules In Vivo. Cell 165, 976–989. 10.1016/j.cell.2016.04.034.

Yasuma, K., Yasunaga, J., Takemoto, K., Sugata, K., Mitobe, Y., Takenouchi, N., Nakagawa, M., Suzuki, Y., and Matsuoka, M. (2016). HTLV-1 bZIP Factor Impairs Anti-viral Immunity by Inducing Co-inhibitory Molecule, T Cell Immunoglobulin and ITIM Domain (TIGIT). PLoS Pathog 12, e1005372. 10.1371/journal.ppat.1005372.

Yoshida, N., Shigemori, K., Donaldson, N., Trevisani, C., Cordero, N.A., Stevenson, K.E., Bu, X., Arakawa, F., Takeuchi, M., Ohshima, K., et al. (2020). Genomic landscape of young ATLL patients identifies frequent targetable CD28 fusions. Blood 135, 1467–1471. 10.1182/blood.2019001815.

Zhang, Y., Liu, T., Meyer, C.A., Eeckhoute, J., Johnson, D.S., Bernstein, B.E., Nusbaum, C., Myers, R.M., Brown, M., Li, W., and Liu, X.S. (2008). Model-based analysis of ChIP-Seq (MACS). Genome Biol 9, R137. 10.1186/gb-2008-9-9-r137.

Zhao, T., Satou, Y., Sugata, K., Miyazato, P., Green, P.L., Imamura, T., and Matsuoka, M. (2011). HTLV-1 bZIP factor enhances TGF-β signaling through p300 coactivator. Blood 118, 1865–1876. 10.1182/blood-2010-12-326199.

Zou, Z., Ohta, T., Miura, F., and Oki, S. (2022). ChIP-Atlas 2021 update: a data-mining suite for exploring epigenomic landscapes by fully integrating ChIP-seq, ATAC-seq and Bisulfite-seq data. Nucleic Acids Research 50, W175–W182. 10.1093/nar/gkac199.

