## Supplemental Figures for "Reprogramming of lactate metabolism is linked to the oncogenesis of the virus-induced leukemia"

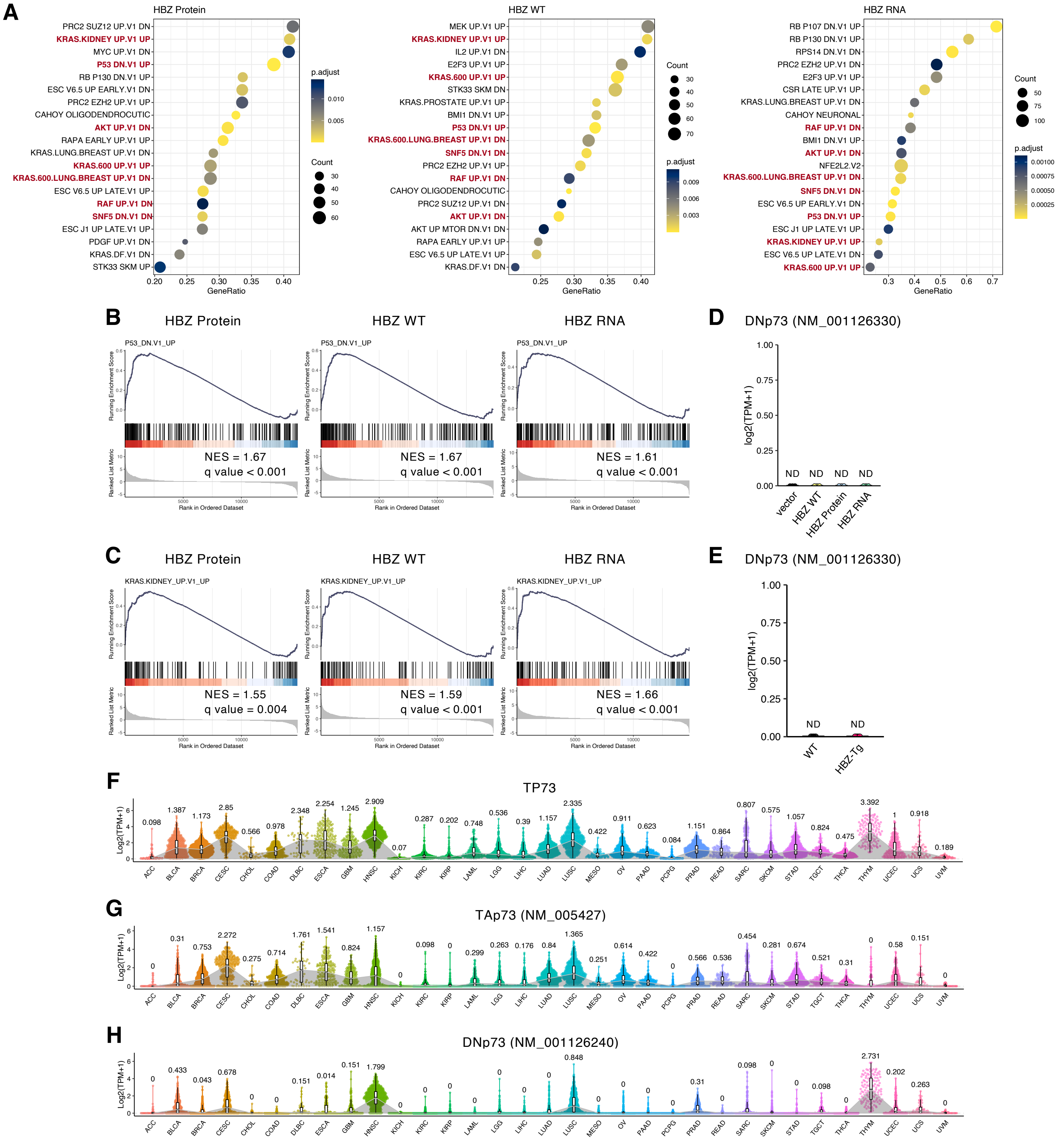


**Figure S1. Transcriptional regulation by HBZ protein and RNA and the status of TP73 expression, related to Figure 1.**

(A) Dot plots of GSEA results among HBZ-transduced murine CD4+ T cells. Statistical values and gene counts determined by the clusterProfiler are shown.
(B and C) Representative GSEA plots, P53_DN.V1_UP (B) and KRAS.KIDNEY_UP.V1_UP (C) for cells transduced with HBZ compared to the vector. The NES and FDR q-values are listed.
(D and E) TPMs of DNp73 in HBZ-transduced murine CD4+ T cells (D); and in WT and HBZ-Tg mice (E). ND, not detectable.
(F-H) TPMs of TP73 (F), TAp73 (G) and DNp73 in TCGA data. Tumor type abbreviations are found in the STAR Methods.

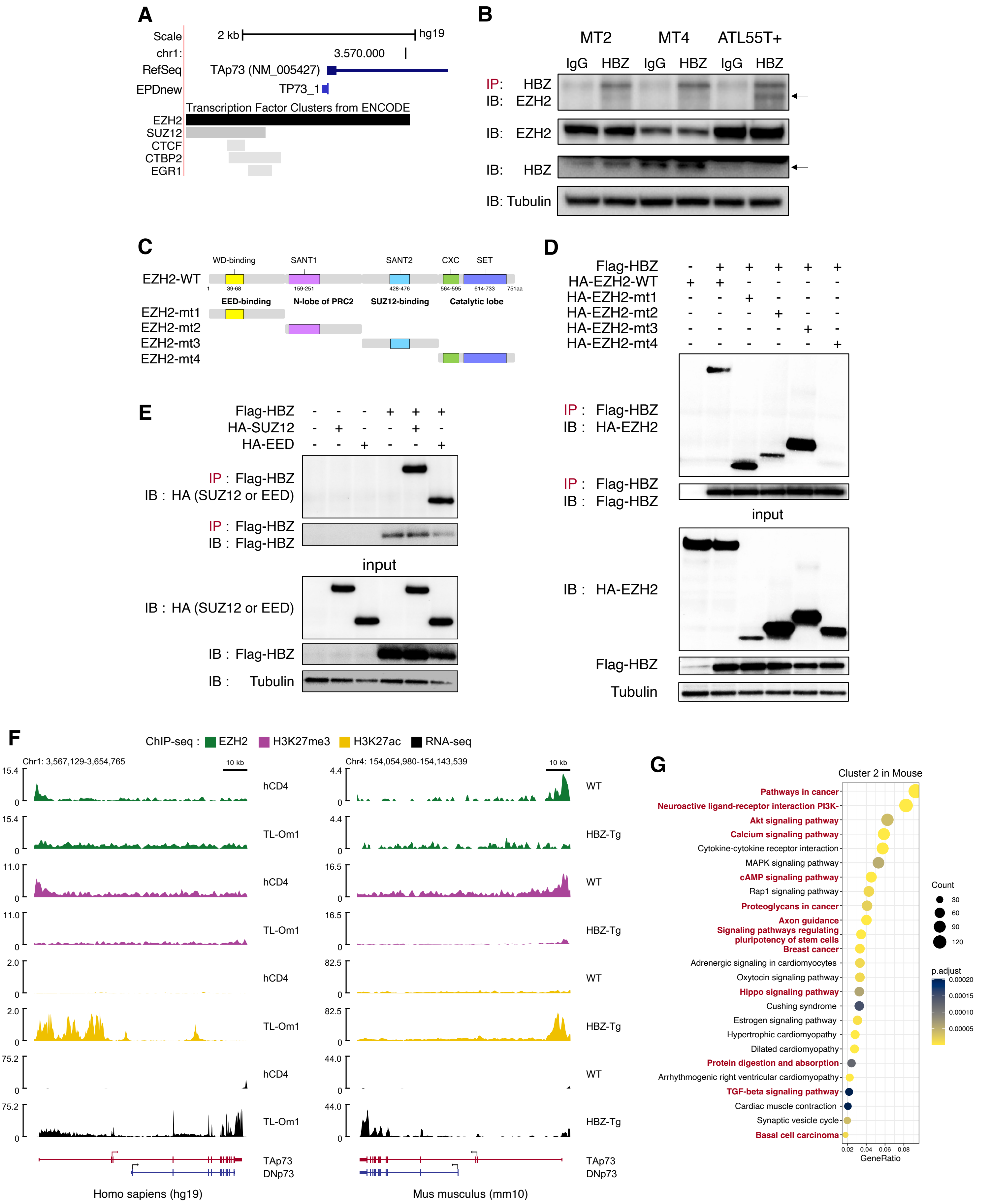

A
Scale
2 kb
hg19
chr1:
3.570.000
RefSeq
TAp73 (NM_005427)
EPDnew
TP73_1
Transcription Factor Clusters from ENCODE
EZH2
SUZ12
CTCF
CTBP2
EGR1
B
MT2
MT4
ATL55T+
IgG
HBZ
IP: HBZ
IB: EZH2
IB: EZH2
IB: HBZ
IB: Tubulin
C
WD-binding
SANT1
SANT2
CXC
SET
EZH2-WT
EED-binding
N-lobe of PRC2
SUZ12-binding
Catalytic lobe
EZH2-mt1
EZH2-mt2
EZH2-mt3
EZH2-mt4
D
Flag-HBZ
HA-EZH2-WT
HA-EZH2-mt1
HA-EZH2-mt2
HA-EZH2-mt3
HA-EZH2-mt4
IP : Flag-HBZ
IB : HA-EZH2
IP : Flag-HBZ
IB : Flag-HBZ
input
IB : HA-EZH2
Flag-HBZ
Tubulin
E
Flag-HBZ
HA-SUZ12
HA-EED
IP : Flag-HBZ
IB : HA (SUZ12 or EED)
IP : Flag-HBZ
IB : Flag-HBZ
input
IB : HA (SUZ12 or EED)
IB : Flag-HBZ
IB : Tubulin
F
ChIP-seq : EZH2 H3K27me3 H3K27ac RNA-seq
Chr1: 3,567,129-3,654,765
10 kb
Chr4: 154,054,980-154,143,539
hCD4
TL-Om1
WT
HBZ-Tg
TAp73
DNp73
Homo sapiens (hg19)
Mus musculus (mm10)
G
Cluster 2 in Mouse
Pathways in cancer
Neuroactive ligand-receptor interaction PI3K-
Akt signaling pathway
Calcium signaling pathway
Cytokine-cytokine receptor interaction
MAPK signaling pathway
cAMP signaling pathway
Rap1 signaling pathway
Proteoglycans in cancer
Axon guidance
Signaling pathways regulating pluripotency of stem cells
Breast cancer
Adrenergic signaling in cardiomyocytes
Oxytocin signaling pathway
Hippo signaling pathway
Cushing syndrome
Estrogen signaling pathway
Hypertrophic cardiomyopathy
Dilated cardiomyopathy
Protein digestion and absorption
Arrhythmogenic right ventricular cardiomyopathy
TGF-beta signaling pathway
Cardiac muscle contraction
Synaptic vesicle cycle
Basal cell carcinoma
GeneRatio
Count
p.adjust

**Figure S2. HBZ protein alters epigenetic regulation of TAp73 by EZH2, related to Figure 2.**

(A) A snapshot of the UCSC genome browser around the TAp73 promoter (registered as TP73_1 in the EPDnew [Eukaryotic Promoter Database]) in hg19. Enriched transcription factors from the ENCODE are shown as barplots.
(B) Endogenous immunoprecipitation (IP) of HBZ protein in HTLV-1-infected cells. IP was performed with anti-HBZ antibody and analyzed by SDS-PAGE and immunoblotting (IB).
(C and D) IP to examine binding cites of HBZ protein to EZH2. A schematic diagram showing the EZH2 mutants and the domains of the EZH2 protein (C). IP of WT or mutant EZH2 with the HBZ protein (anti-Flag antibody) in 293T cells (D).
(E) IP of SUZ12 or EED with the HBZ protein (anti-Flag antibody) in 293T cells.
(F) EZH2, H3K27me3 and H3K27ac enrichments (ChIP-seq) and transcripts (RNA-seq) around Trp73 in WT or HBZ-Tg mouse $CD4^+$ T cells (left), and around TP73 in hCD4 and TL-Om1 cells (right).
(G) Results of KEGG pathway analysis using the mouse cluster 2 genes (Figure 2H). Statistical values and gene counts calculated by the clusterProfiler are shown.

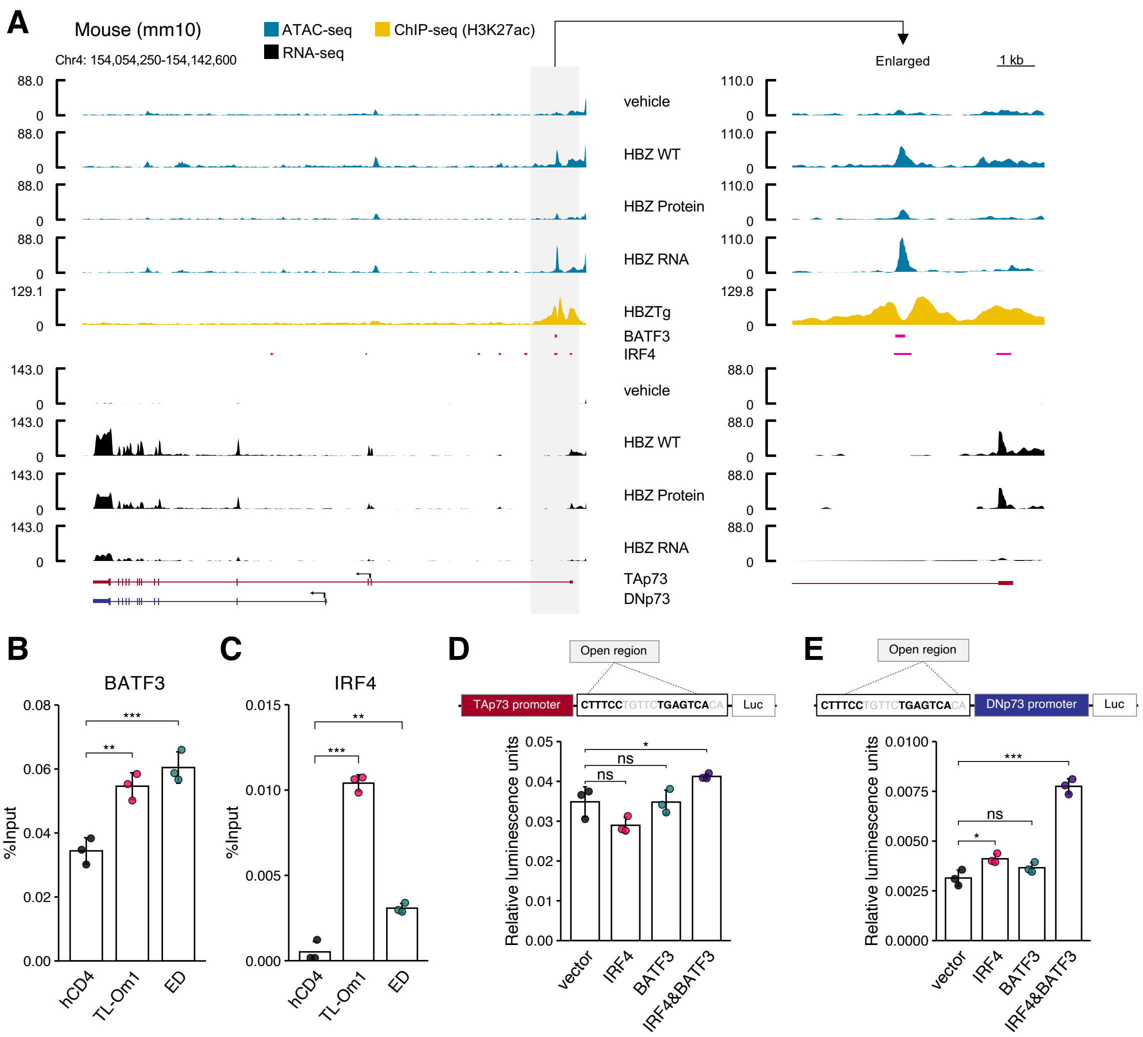


**Figure S3. Regulation of TP73 by BATF3 and IRF4, related to Figure 3.**

(A) H3K27ac enrichment, BATF3/IRF4 binding of ENCODE data (ChIP-seq; SRX2646182 and SRX2646184) (Iwata *et al.*, 2017), Chromatin accessibility (ATAC-seq) and transcription (RNA-seq) of Trp73 in HBZ-transduced murine CD4+ T cells.
(B and C) ChIP-qPCR in ATL cells lines for BATF3 (B) and IRF4 (C) in the open region found in Figure 3C (n=3).
(D and E) Promoter assays of the open region in Jurkat cells. The IRF4/AP-1 motifs identified within the open region were subjected to promoter assays with IRF4 and/or BATF3 induction for the promoters of TAp73 (D) and DNp73 (E) (n=3). A schematic of the assay construct is shown above the corresponding bar plot.
Results are plotted as mean ± SD, using one-way ANOVA with Dunnet (B-E). $*p < 0.05$, $**p < 0.01$, $***p < 0.001$; ns, not significant.

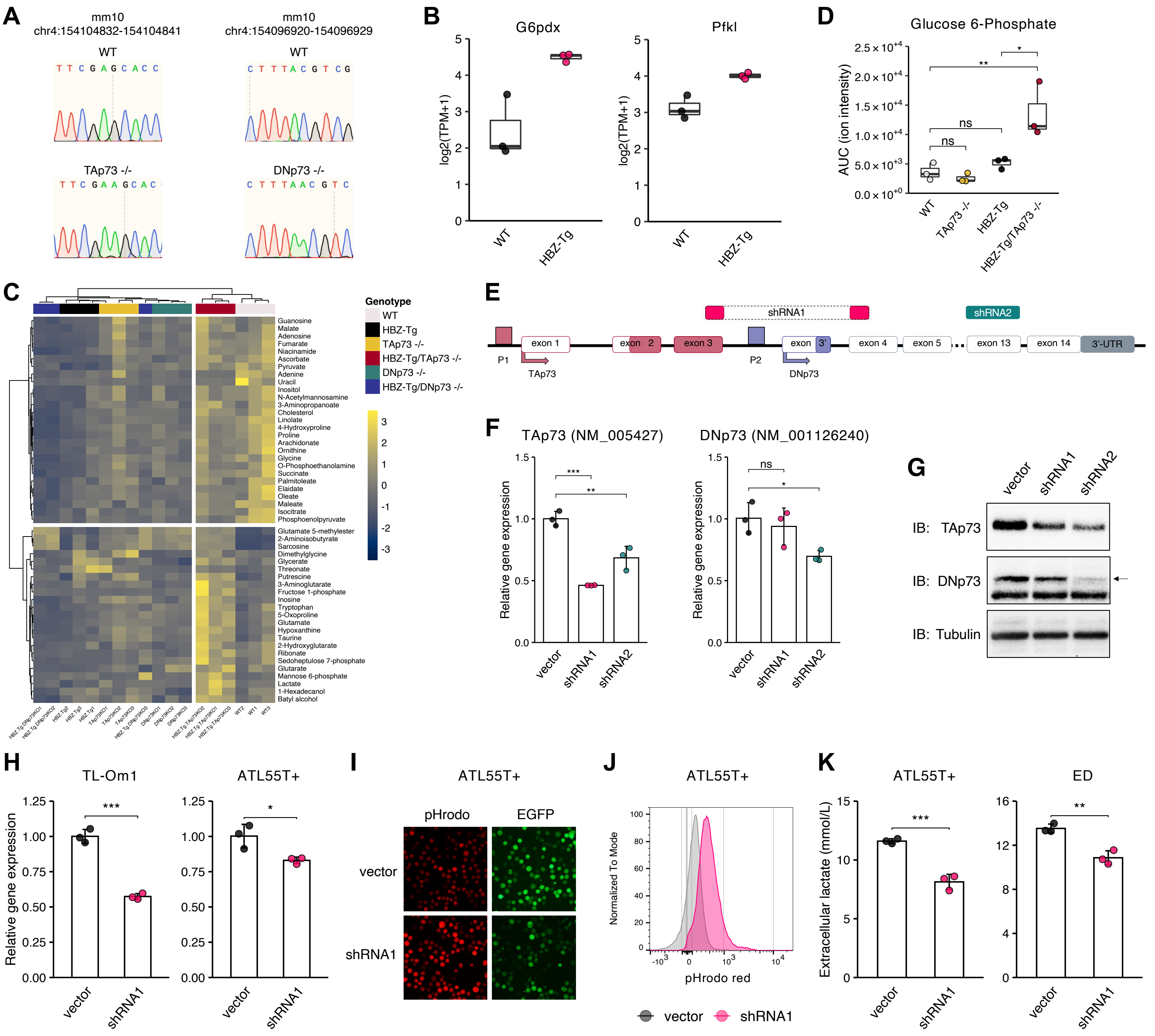

**Figure S4. Transcriptional regulation of SLC16A1 and SLC16A3 by TAp73 and mRNA expression levels of MCT-coding genes among various cancers, related to Figure 5.**

(A) Sequencing results of Trp73-isoform specific murine KO models. Each 1bp frameshift resulted in a truncating mutation.
(B) TPMs of G6odx (left) and Pfkl (right) in WT or HBZ-Tg mouse CD4⁺ T cells.
(C) A heatmap of the top 50 differentially altered metabolites among the KO murine models.
(D) Glucose 6-Phosphate levels in CD4⁺ T cells of the murine models (n=3). Calculated ion intensities (AUC) are shown.
(E-G) KD of the human TP73 gene. Schematic diagram showing two kinds of shRNAs for the human TP73 gene (E). mRNA expressions of TAp73 (left) and DNp73 (right) by RT-qPCR in TL-Om1 cells 48 hours after the KD (F) (n=3). Immunoblots of TAp73, DNp73 and Tubulin in 293T cells with ectopic expression of shRNA (G).
(H) EZH2 transcripts in ATL cells 48 hours after the KD (n=3).
(I and J) Intracellular pH assessed by pHrodo Red AM in ATL55T+ cells on day 8 after transfection. Fluorescence microscopy photographs with EGFP (I) and a flow cytometry histogram (J).
(K) Extracellular lactate from ATL cells on day 8 after transfection (n=3).
Results are plotted as mean ± SD, using one-way ANOVA with post-hoc Turkey (D), Dunnet (F) or Student's t test (H and K). *p < 0.05, **p < 0.01, ***p < 0.001; ns, not significant.

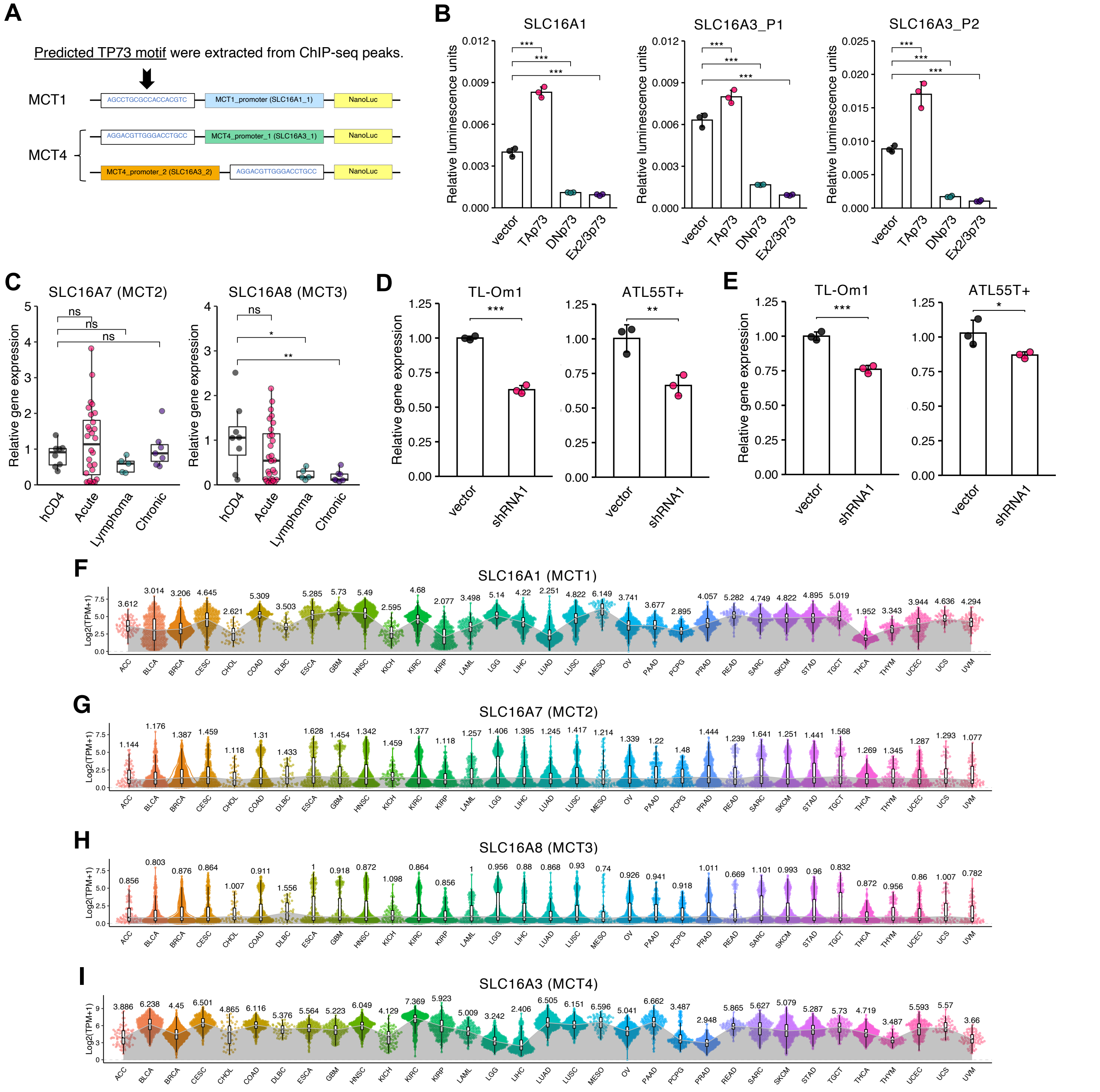


**Figure S5. Transcriptional regulation of SLC16A1 and SLC16A3 by TAp73, and mRNA expression levels of MCT-coding genes among various cancers, related to Figure 6.**

(A) A schematic of vectors for the luciferase assay related to Figures 6C and S5B. Enriched TP73 binding sequences from the ChIP-seq peaks for TAp73 are inserted into the vectors with SLC16A1 and SLC16A3 promoters. Since two kinds of promoters have been reported for the latter, we generated two corresponding vectors as shown.
(B) Relative luciferase activities for the SLC16A1 promoter (left) and the SLC16A3 promoter (center and right) with TP73 isoform expression in Jurkat cells. (n=3).
(C) mRNA expression of SLC16A7 (left) and SLC16A8 (right) by RT-qPCR in hCD4 (n=6) and ATL cells (acute type, n=32; lymphoma type, n=7; chronic type, n=7).
(D and E) mRNA transcripts of SLC16A1 (D) and SLC16A3 (E) in ATL cells after TAp73 KD for 48 hours (n=3).
(F-I) TPMs of MCT family-coding genes in TCGA data. Tumor type abbreviations are found in the STAR Methods.
Results are plotted as mean ± SD, using the Student's t test (D and E), one-way ANOVA with post-hoc Dunnet (B) or Steel test (C). *p < 0.05, **p < 0.01, ***p < 0.001; ns, not significant.

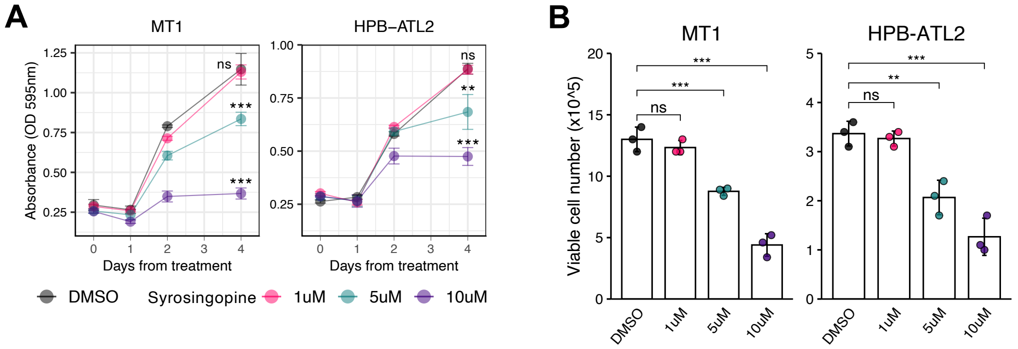


**Figure S6. Efficacy of the MCT1/4 inhibitor syrosingopine on ATL cells, related to Figure 7.**

(A and B) Cell proliferation assay (A) and viable cell numbers (B) for ATL cells (MT1 and HPB-ATL2) treated with syrosingopine (1μM, 5μM and 10μM; n=3). Values in comparison to the DMSO group are shown.
Results are plotted as mean ± SD, using one-way ANOVA with post-hoc Dunnet test (A and B). **$p < 0.01$, ***$p < 0.001$; ns, not significant.
